# MEBRAINS 1.0: a new population-based macaque atlas

**DOI:** 10.1101/2023.06.21.545953

**Authors:** Puiu F Balan, Qi Zhu, Xiaolian Li, Meiqi Niu, Lucija Rapan, Thomas Funck, Rembrandt Bakker, Nicola Palomero-Gallagher, Wim Vanduffel

**Author notes:** **corresponding authors:** Wim Vanduffel; Nicola Palomero-Gallagher. Authors contributed equally (Puiu F Balan, Qi Zhu). Joint senior authorship (Nicola Palomero-Gallagher, Wim Vanduffel).

## Abstract

Due to their fundamental relevance, the number of anatomical macaque brain templates is constantly growing. Novel templates aim to alleviate limitations of previously published atlases and offer the foundation to integrate multiscale multimodal data. Typical limitations of existing templates include their reliance on one subject, their unimodality (usually only T1 or histological images), or lack of anatomical details. The MEBRAINS template overcomes these limitations by using a combination of T1 and T2 images, from the same 10 animals (Macaca mulatta), which are averaged by the multi-brain toolbox for diffeomorphic registration and segmentation. The resulting volumetric T1 and T2 templates are supplemented with high quality white and gray matter surfaces built with FreeSurfer. Human-curated segmentations of pial surface, white/gray matter interface and major subcortical nuclei were used to analyse the relative quality of the MEBRAINS template. Recently published 3D maps of the macaque inferior parietal lobe and (pre)motor cortex were warped to the MEBRAINS surface template, thus populating it with a parcellation scheme based on cyto- and receptor architectonic analyses. Finally, 9 CT scans of the same monkeys were registered to the T1 modality and co-registered to the template. Through its main features (multi-subject, multi-modal, volume-and-surface, traditional and deep learning-based segmentations), MEBRAINS aims to improve integration of multi-modal multi-scale macaque data and is quantitatively equal or better compared to currently widely used macaque templates. The template is integrated in the EBRAINS and Scalable Brain Atlas web-based infrastructures, each of which comes with its own suite of spatial registration tools.

## INTRODUCTION

The macaque monkey is an important model system for systems neuroscience. Genetic, functional, and anatomical properties of the macaque brain resemble those of the human more closely than other animal models which can be used in biomedical research. As such the macaque has provided translational benefits and the ability to test hypotheses using very precise invasive techniques (e.g., electrophysiology, optogenetics, histology, lesions, etc.). Moreover, the application of non-invasive brain imaging techniques in both humans and monkeys has helped to relate hemodynamic findings from human research to neuronal properties and demonstrate the translational relevance of the macaque as a model system (Seidlitz et al., 2018).

The existence of anatomical templates is an essential step, however, to anchor and integrate a wealth of multi-level neuroscience data (from molecules to maps) in the same ordered space and to enable objective cross-level or cross-species comparisons, an approach which has recently been implemented for the human brain (Amunts et al., 2014). Single subject-based neuroscience is by definition limited by the idiosyncratic anatomy and physiology of an individual, hence does not allow us to make general statements at population level. Multi-subject analyses, on the other hand, bolster scientific validity by increasing statistical power and highlighting reliable neurological phenomena across the population (Friston et al., 1999). To facilitate comparisons across subjects, data from each subject should be registered to a template. Moreover, templates based on multiple subjects are optimal for group-level analyses because they possess features that are more representative of the population’s “average” brain anatomy which offers higher cross-subject validity (Dadar et al., 2022; Evans et al., 2012; Fonov et al., 2011).

Because of their value, macaque neuroscience is populated with increasingly more and better anatomical templates (Table 1), each with their own benefits and caveats. Fortunately, mathematical transformations allow us to link representations between different template spaces. In line with this, also the number of publications (Figure 1) related to research using macaque brain templates is increasing.

**Figure 1.**
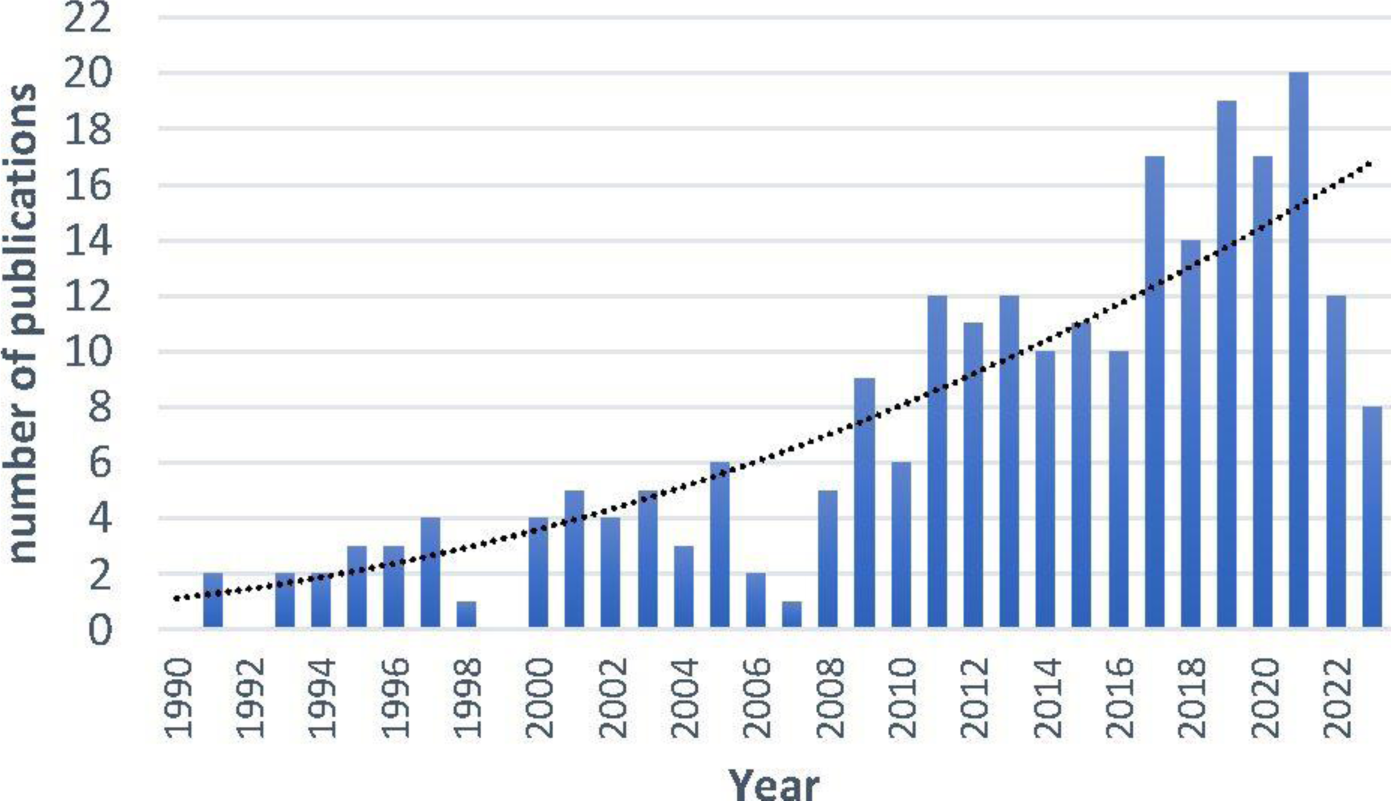
Number of publications per year related to brain templates in macaque monkeys. A PubMed search query was performed June 2023 using the following keyword combination: (“monkey” OR “macaque” OR “NHP” OR “non-human primate”) AND (“template” OR “atlas”) AND (brain). Polynomial fit with R^2^ = 0.7157.

**Table 1.** Non-exhaustive list of some of the most frequently used macaque templates. All templates were obtained from *Macaca mulatta* monkeys, except for the MNI template, which was built from *Macaca mulatta* (Mm) and *Macaca fascicularis* (Mf) brain scans. Abbreviations: N/A = not available; Res. = Resolution; Skull str. = the template is available in the original format (OF) or only in a skull stripped (SSF) format.

| Template | Skull str. | Sequence | Res. (mm) | Number of brains | Associated atlas(es) | Volume format | Surface format |
| --- | --- | --- | --- | --- | --- | --- | --- |
| NMT (Jung et al., 2021; Seidlitz et al., 2018) v1.2/v1.3/v2.0 | OF | T1 | 0.25 | 31 | D99-SL (Reveley et al., 2017)<br>CHARM (Jung et al., 2021)<br>SARM (Hartig et al., 2021) | NIFTI | GIFTI |
| D99 (Reveley et al., 2017; Saleem et al., 2021) v1/v2 | SSF | T1, T2, DTI, MAP-MRI, MTR | 0.25 | 1 | D99-SL | NIFTI | GIFTI |
| INIA19 (Rohlfing et al., 2012) | OF | T1 | 0.50 | 19 | Neuromaps | NIFTI | N/A |
| MNI (Frey et al., 2011) | OF | T1 | 0.25 | 18 Mf<br>7Mm | Paxinos | MINC, NIFTI | N/A |
| Yerkes19 (Donahue et al., 2018; Van Essen et al., 2012) | OF | T1, T2 | 0.50 | 19 | F99 (Van Essen, 2004) | NIFTI, MGZ | GIFTI, MGZ |
| 112RM-SL (McLaren et al., 2009) | SSF | T1, T2* | 0.50 | 112 (McLaren et al., 2009)* | D99-SL (Reveley et al., 2017)<br>F99 (Van Essen, 2004) | NIFTI | N/A |
| UNC-Emory atlas (Shi et al., 2016) | OF | T1, T2, DTI | 0.60 | 40 |  | NRRD | N/A |
| ONPRC18 (Weiss et al., 2021) | SSF | T1, T2, DTI | 0.50 | 18 | ONPRC18 | NIFTI | N/A |
| F99 (Van Essen, 2004) | SSF | T1 | 0.50 | 1 |  | NIFTI | GIFTI |

\* T2-weighted scans only available for 9 of the 112 animals

However, existing templates have important limitations when they are based on a single animal, unimodal images (e.g., T1-weighted images), or when they lack sufficient anatomical details (i.e., when the resolution is too low). While single subject-based templates are less representative of the population’s anatomy, multi-subject templates suffer from blurred images because of non-perfect registration between images of the individual subjects and inherent averaging-induced smoothing. Recently, multi-subject templates have been improved relative to those which were based on linear registration methods (Friston et al., 1999) by employing sophisticated nonlinear transformation techniques (Brudfors et al., 2020; Friston et al., 1999). These novel methods (Brudfors et al., 2020) yielded improved anatomical details and contrast. However, nonlinear transformation algorithms on 3D volumes easily result in warping artefacts due to their high degrees of freedom and flexibility. Consequently, there is a strong interest to use surfaces for displaying data and registering brain images. Yet, multi-subject templates providing surfaces in addition to volumetric representation are still rare (see Table 1).

To address this problem, we propose a first version of a template based on the brains of 10 monkeys for which both high-resolution (isotropic 0.064 mm^3^) T1 and T2 images were recorded within the same scan session. Additionally, CT scans are available for 9 of these monkeys. We are steadily increasing the number of subjects, which will be implemented in later versions of the template. Second, we tested and compared several non-linear registration algorithms to improve the quality of the average template. The multi-brain (MB) toolbox (Brudfors et al., 2020) applied simultaneously to T1 and T2 images resulted in the most faithful template and was selected as the best solution. Additionally, it generates an underlying tissue classification as part of the registration process. Third, our approach allows to integrate an unlimited number of modalities (e.g., T1, T2, diffusion-weighted (DW), computed tomography (CT)) using the same processing software. Fourth, we provide both volumetric and surface representations of the template. Fifth, our template is integrated in the EBRAINS environment (https://ebrains.eu/about) and thus enables to compare data from multiple species using the same meta-platform. Sixth, we started to populate the template with a human-curated segmentation of major subcortical nuclei and with recently published maps of the macaque monkey motor, parietal and early visual cortex based on cyto- and receptor architectonic analyses (Niu et al., 2020; Niu et al., 2021; Rapan et al., 2021; Rapan et al., 2022). Seventh, we integrated new methods for data processing in the macaque based on recent AI developments and applications in neuroscience, (e.g., deep learning for skull stripping and segmentation). Last, but not least, several of the animals with brain anatomies included in this template are still alive, so new data can be acquired to populate and enrich the atlas.

## MATERIALS AND METHODS

### Subject information

10 rhesus monkeys (Macaca mulatta; 3 female) were used in this study. The monkeys were young adults, with an average age of 5.30 year (6.33 for female, and 4.86 for male) when the anatomical scans were collected. The monkeys weighted 6.33 kg on average (5.50 kg for the females, and 8.00 kg for the males) at the time of scanning. Animal care and experimental procedures were performed in accordance with the National Institute of Health’s Guide for the Care and Use of Laboratory Animal, the European legislation (Directive 2010/63/EU) and were approved by the Animal Ethics Committee of the KU Leuven. Weatherall reports were used as reference for animal housing and handling. All animals were group-housed in cages sized 16-32 m^3^, which encourages social interactions and locomotor behavior. The environment was enriched by foraging devices and toys. The animals were fed daily with standard primate chow supplemented with fruits, vegetables, bread, peanuts, cashew nuts, raisins and dry apricots. They had free water access during the period that the anatomical scans were acquired. All animals participated in behavioral, fMRI, electrophysiology and/or reversible perturbation experiments afterwards (Arsenault et al., 2014; Arsenault and Vanduffel, 2019; Balan et al., 2018; Caspari et al., 2015; Herpers et al., 2021; Janssens et al., 2014; Li et al., 2022; Murris et al., 2021; Yao and Vanduffel, 2022).

### Acquisition of anatomical MR and CT images

High-resolution (400 μm isotropic voxel size) T1- and T2-weighted images were acquired on a 3T Siemens PrismaFit scanner while the animals were under ketamine/xylazine anaesthesia. A custom-built single loop coil with a diameter of 12 cm was used as receiver, and the body coil from the scanner was used for transmission. T1 images were acquired using a magnetization prepared rapid gradient echo (MPRAGE) sequence (repetition time (TR) = 2700 ms, echo time (TE) = 3.5 ms, flip angle (α) = 9°, inversion time (TI) = 882 ms, matrix size 320×260×208) and T2 images were acquired using a sampling perfection with application optimized contrasts using different flip angle evolution (SPACE) sequence (TR = 3200 ms, TE = 456 ms, variable α, matrix size 320 × 260 × 208, Turbo Factor = 131, echo spacing = 6 ms), as in (Glasser and Essen, 2011; Van Essen et al., 2001). During a single scan session, 7–12 T1 images and 4–5 T2 images were acquired from each subject (Li et al., 2021). Additionally, for 9 of the animals, high resolution CT (324x324x200 matrix size; 0.25 mm isotropic; on a Somatom Force Siemens CT scanner) scans were acquired in different sessions while the animals were under ketamine/xylazine anaesthesia.

Pre-processing of these images for their compatibility with Freesurfer and MB constituted the first step of the pipeline developed for the development of the template (Figure 2).

**Figure 2.**
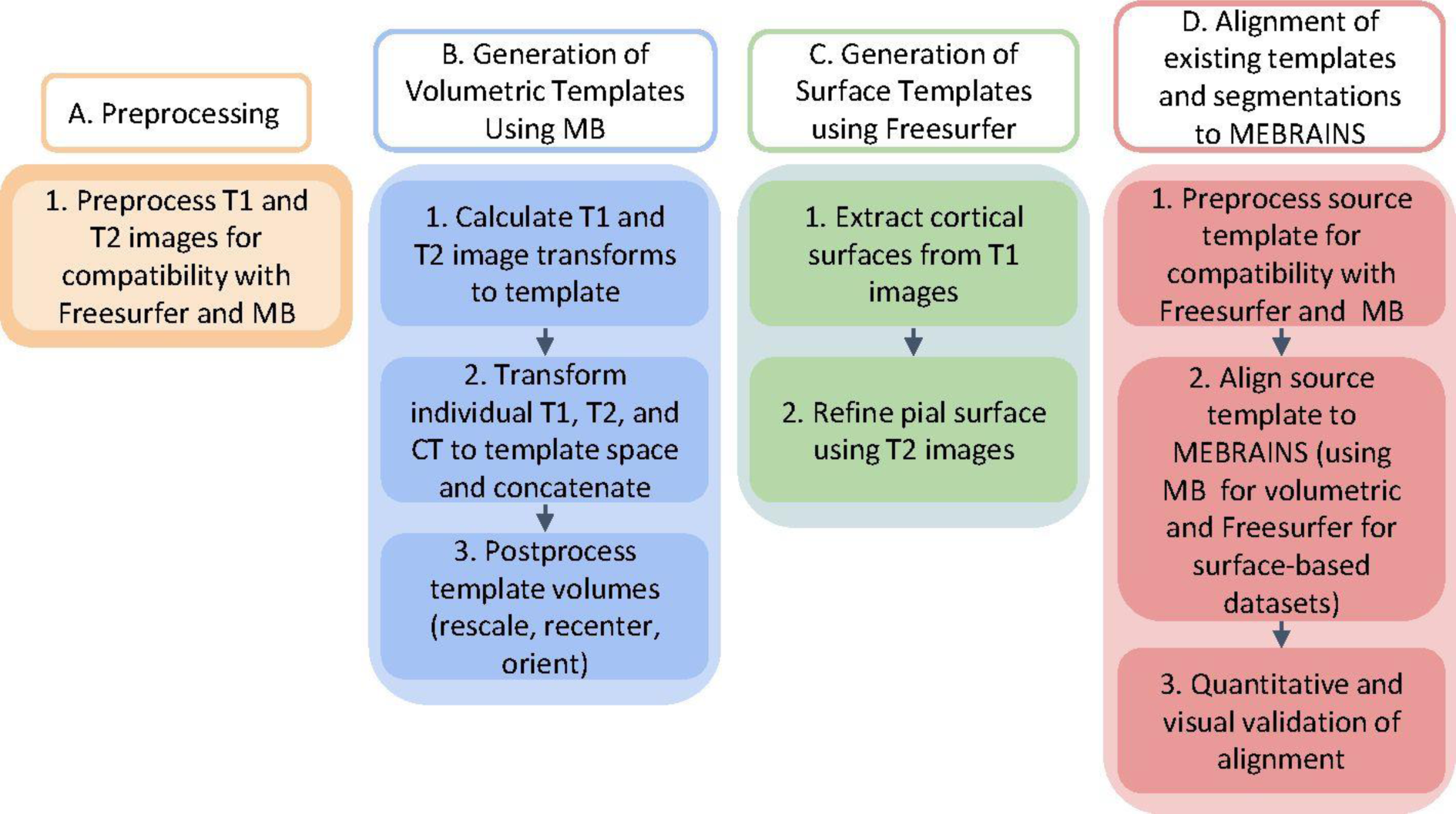
Overview of the pipeline used for the generation of a population-based template that represents an average of high-resolution structural T1 and T2 MRI scans as well as CT.

### Anatomical MR and CT pre-processing (Autio et al., 2021)

The pre-processing consisted of:

- DICOM to NIFTI conversion of both MR and CT datasets using FreeSurfer (Fischl, 2012).
- Per subject, registration of the CT to the corresponding anatomical MR using FreeSurfer, ANTS (Avants et al., 2011), and ITK-SNAP (Yushkevich et al., 2006).
- Conversion of all volumes to the FreeSurfer-conform standard (256x256x256, orientation LIA (left-inferior-anterior)). The FreeSurfer-conform standard requires 1 mm isotropic voxel size. To satisfy this condition without losing resolution, we arbitrarily changed the voxel size in the image header from 0.4 to 1 mm.
- Rigid registration of all T1 volumes to a unique template (which was the average of all individual T1 volumes which were registered using a pre-run of the multi-brain (MB) toolbox for SPM12 on the original T1 volumes) using a combination of FreeSurfer, ANTS and the MB toolbox. T1, T2 and CT volumes were registered using unique transformation matrices (generated when the T1 volumes were registered) for each subject.
- Bias field correction of the MR anatomies following the Human Connectome Protocol adapted to the macaque (Autio et al., 2021; Hayashi et al., 2021; Marcus et al., 2013).
- To generate symmetrical templates, we added to the existing set of volumes (separately for T1, T2 and CT) their left-right flipped version generated using FreeSurfer.

### Generation of the volumetric anatomical templates using T1 and T2 anatomies

#### MEBRAINS template construction with the multi-brain toolbox

The main processing tool for building the MEBRAINS template was the MB toolbox of SPM12 (Brudfors et al., 2020) (https://github.com/WTCN-computational-anatomy-group/mb), and as input we used information from both T1 and T2 images. We chose MB because it generates a probabilistic tissue classification model while performing the nonlinear registration, rather than just using voxel intensities directly. This approach has been shown to be a more robust method of registering medical images (Klein et al., 2009; Sotiras et al., 2013). Furthermore, the algorithm (Brudfors et al., 2020) used by MB can integrate many imaging modalities (e.g., T1, T2, DW, CT), and can be applied with or without prior pre-processing (e.g., skull stripping). Accordingly, we took advantage of the high-resolution CT scans of the same subjects, applied the same transformations as those used to register the corresponding T1 and T2 images to the reference template, and averaged the resulting CTs to build the CT template. Thus, multi-brain allowed us to build the following three templates using T1, T2 and CT brain images of 10 monkeys: MEBRAINS_T1, MEBRAINS_T2 and MEBRAINS_CT, respectively. We generated the volumetric templates as follows:

i. Learn the MB tissue probability model. We adapted Example 1 from the MB repository (https://github.com/WTCN-computational-anatomy-group/mb). As input we used the set of 10 pairs of T1 and T2 images and additionally the same set of images mirrored across the midsagittal plane to create a symmetric template. This group-wise image registration generated the following datasets: an optimal K class tissue template; optimal intensity parameters; deformations that are used to warp between different volumes; tissue segmentations; and bias-field corrected versions of the input scans. In general, we kept the default settings to run the MB modelling (as in Example 1 mentioned above). The following parameters were modified in our script: regularization of the nonlinear registration (changed from 1 to 2), number of tissue types K (set to 14), and voxel size (set to 1).
ii. Register the T1 and T2 individual volumes to the MB tissue model using the MB deformations generated during the learning step, as in example 2 of the MB repository (https://github.com/WTCN-computational-anatomy-group/mb). We used a 3^rd^ degree B-spline interpolation algorithm, and co-registered the CT volumes with the T1 volumes.
iii. Create T1, T2 and CT templates by averaging the corresponding individual images registered to the MB tissue model. Intermediate T1, T2 and CT templates are created by gradually averaging more and more individual images that are registered to the implicit MB template.
iv. Linear transformation of the templates to set each origin to the center of the anterior commissure as identified in a sagittal section (voxel 108,128,70 in RAS-coordinates, i.e., with voxel 0,0,0 at the left-posterior-inferior corner).
v. Rescale the volumes to the original resolution of 0.4 mm isotropic voxels.
vi. Check the stereotaxic orientation of the template. Since the original brains were acquired using a stereotaxic frame, we verified that the resulting average has the aural fixation points and the infraorbital ridge nearly in the same horizontal plane, which is a requirement of being aligned to the Horsley-Clarke stereotaxic frame (Seidlitz et al., 2018).

#### Comparative template – ANTS10

The ANTS version of the template was built as a comparison with MB in terms of warping artefacts. We followed the processing described in (Seidlitz et al., 2018) and used whole-head images so that the template would accurately represent the brain-skull boundary. The main processing steps were:

i. Align each of the 10 preprocessed subject images to an independent coordinate space (EBRAINS_T1) using a 6-parameter rigid-body transformation.
ii. Create the initial target image for the template by performing a voxel-wise average of the 10 subject images.
iii. Normalization of the variations in image intensity across each volume by an N4 bias field correction (Avants et al., 2011).
iv. Create the population-averaged template using symmetrical group-wise normalization, which is an iterative nonlinear registration process (Seidlitz et al., 2018). Each brain was aligned to the current target image via a 12-parameter affine and a nonlinear (diffeomorphic) transformation. These aligned images were averaged to generate an improved template image. The inverse of the affine and diffeomorphic transformations was averaged across subjects, scaled, and applied to this template image to align it closer to the original input anatomies. This process was iterated, with the updated template image serving as the new target image for registration with the original subject images, until convergence between successive target images occurred.

### Generation of a MEBRAINS surface template

Surface representations of the brain enable a more precise spatial localization and reduce the occurrence of errors arising from the spatial proximity of brain structures that are actually located at quite a distance from each other along the cortical ribbon (Logothetis et al., 2001; Zhu and Vanduffel, 2019). Additionally, they are a prerequisite for generating cortical flat maps, which are useful tools for the analysis and visualization of functional and structural neuroimaging datasets (Sultan et al., 2010; Van Essen et al., 1998; Vanduffel et al., 2001; Vanduffel et al., 2014), particularly for topographic representations such as retinotopy (Arcaro and Livingstone, 2017; Janssens et al., 2014), somatotopy (Arcaro et al., 2019) and tonotopy (Bodin et al., 2021; Erb et al., 2019; Petkov et al., 2006). To achieve this, a human-curated white and gray matter segmentation was performed with FreeSurfer (Fischl, 2012) and the non-human primate version of the Human Connectome Project pipeline (Autio et al., 2020), using a combination of T1 and T2 images (Autio et al., 2021). The pial and white/gray matter interface (white matter surface) was generated from the T1 images to create the MEBRAINS surface template. T2 images were used to accurately model the pial surface and remove the effect of cerebrospinal fluid and pial veins.

### “Populating” the MEBRAINS template: human-curated segmentations of subcortical nuclei and integration of cyto- and receptor architectonically informed cortical maps

We started to populate the template by complementing MEBRAINS with human-curated segmentations of several subcortical structures. We manually delineated the amygdala, anterior commissure, nucleus accumbens, caudate, claustrum, putamen, and pallidum on coronal sections of the left hemisphere of the MEBRAINS_T1 template, whereby all three stereotactic planes were closely examined to reduce inconsistencies across slices. This segmentation was performed using MRIcron (Rorden and Brett, 2000) and ITKsnap (Yushkevich et al., 2006), and identification of structures was based on local contrast differences in both the EBRAINS_T1 and the EBRAINS_T2 templates, thereby relying on corresponding sections of the 2^nd^ edition of the Atlas of the Rhesus Monkey Brain (Saleem and Nikos, 2012). The delineated structures were mirrored (using MATLAB, FreeSurfer and human-curation) to segment the right hemisphere of the template. These human-curated segmentations were also essential for our quality assessment of MEBRAINS and to develop workflows for integrating 3D volumes into MEBRAINS space. Specifically, these segmentations i) served as a reference when evaluating the quality of (semi)-automated segmentation approaches, and ii) generated target outputs (ground-truth) for training deep neural networks to automatically segment brain structures (Henschel et al., 2020).

Additionally, we used the workflow to integrate other templates into MEBRAINS, for example, to anchor the frequently used D99-atlas and our recently published 3D cyto- and receptor architectonic maps of the macaque parietal (Impieri et al., 2019; Niu et al., 2020; Niu et al., 2021), premotor and motor (Rapan et al., 2021) cortex, depicted on the Yerkes19 template (Donahue et al., 2018; Van Essen et al., 2012) into MEBRAINS space. Since the MEBRAINS template is symmetrical, and these parcellations were only available for the left hemisphere of the Yerkes template, the ensuing maps had to be human-curated using ITKsnap (Yushkevich et al., 2006), then mirrored to the right hemisphere of MEBRAINS using MATLAB and FreeSurfer.

### Registration of 3D datasets to MEBRAINS

Since it is essential to link MEBRAINS to commonly used template spaces, we developed a multi-method workflow to register 3D data to MEBRAINS. Independent of the method/algorithm used, registration of 3D volumes can be achieved as follows:

- **Step 1.** Preparatory pre-processing of the data to roughly adjust the image geometry (i.e., resolution, dimensions, position) performed with FreeSurfer, FSL (Woolrich et al., 2009) and MATLAB. This step does not necessarily require MB.
- **Step 2.** Register the brain anatomy (e.g., other template volume or individual anatomy) to MEBRAINS. This process is achieved by calculating and applying the transformation functions (matrices and deformation volumes). Noteworthy, the transformations generated for a specific volume (e.g., a template) can be applied to different entities (e.g., atlas, connectivity maps) represented in that space. The specifics of the registration performed with MB are found under “https://github.com/WTCN-computational-anatomy-group/mb - Example 3: Fitting a learned MB model”, and were applied to individual brain anatomy/template volumes.
- **Step 3.** Evaluate the quality of the registration and improve it by adjusting different parameters of the registration algorithm. If the object to be registered is a template brain or an individual anatomical dataset, the process is finished. We used “https://github.com/WTCN-computational-anatomy-group/mb - 2. Warping with MB deformations - image-to-template – pull” to apply the deformation generated in the previous step to the brain anatomy/template.
- **Step 4.** If we register atlases, activation maps, retinotopic maps, or connectivity maps to MEBRAINS, a supplementary step may be necessary because such data require an underlying reference anatomy. This reference anatomy should follow steps 1 to 3, to generate the corresponding transformations/deformations functions to be applied. It is important to remember that resampling algorithms can be nonlinear (e.g., cubic) when transforming anatomical volumes, and resampling algorithms used to register atlases (representing discrete values) should be linear or nearest-neighborhood. The specifics for registrations performed with MB are listed in “https://github.com/WTCN-computational-anatomy-group/mb; “4. Register and warp atlas to MB space<u>“.</u>

Since no single tool functions seamlessly, the best strategy is to combine functions from different software packages. This is illustrated by the existence of an open-source, community-developed initiative like Nypype (Gorgolewski et al., 2011) (https://nipype.readthedocs.io/en/latest/), facilitating interactions between different software packages (e.g., ANTS, SPM, FSL, FreeSurfer, Camino, MRtrix, MNE, AFNI, Slicer, DIPY).

Like all methods, MB also harbors some problems. For example, recall that the MEBRAINS template is built using both T1 and T2 weighted images. If other volumes have to be registered to MEBRAINS, these data contain optimally both T1 and T2 modalities. Furthermore, if we start from already skull-stripped anatomies instead of the whole head, the registration may be sub-optimal.

### A library of registration methods

Although we selected MB as our method of choice to generate the average template, the resulting MEBRAINS template can be used with any registration method. The most relevant software packages are summarized below:

a. Multi-brain (Brudfors et al., 2020) – using MATLAB and toolboxes.
b. ANTS (Avants et al., 2009) – using either the RheMAP (Sirmpilatze and Klink, 2020) Jupiter notebook (https://github.com/PRIME-RE/RheMAP.git), or antsRegistrationSyNQuick to generate the registration and antsApplyTransforms to apply it.
c. AFNI (Cox, 1996) – generate the registration with 3dQwarp and apply it with 3dNwarpApply.
d. MINC (Vincent et al., 2016) – generate the registration with minctracc and apply it with mincresample.
e. ART (Ardekani et al., 2005) – generate the registration with 3dwarper and apply it with applywarp3d.
f. ITKsnap (Yushkevich et al., 2006) – for illustrative affine registrations.
g. FSL (Woolrich et al., 2009) – generate registrations with flirt and fnirt, and apply it with applywarp.
h. Jip (http://www.nitrc.org/projects/jip) – using jip_align in two stages: auto-align affine followed by auto-align non-lin.
i. DISCO (Ardekani et al., 2005) – using the Diffeomorphic Sulcal-based COrtical (DISCO) registration.
j. FreeSurfer (Fischl et al., 1999) – perform either a surface based registration using mris_register, or a combined surface and volume morph method (Postelnicu et al., 2009; Zöllei et al., 2010) using mri_cvs_register. The latter approach accurately registers both cortical and subcortical regions, establishing a single coordinate system suitable for the entire brain.

Many of these tools (a - f) can rapidly register source with target volumes. The others (especially i - j) are computationally costly, and are mainly recommended when the ‘fast’ methods yield suboptimal results.

This library of methods raises a fundamental question: which strategy should one use? We propose the following:

a. Use your own knowledge/preference, but consider the quality of the source anatomy that has to be registered (e.g., template).
b. *Try-N-select-winner*. The strategy works with anatomies and involves the following straightforward steps:

1. Select a registration method and optimize the results by adjusting the parameters of the algorithm.
2. If the result is not satisfactory, add a new method and repeat 1.
3. Compare the existing results and select a winner.
4. If the winner is not satisfactory, repeat 2. If the winner meets your needs stop the process. We list a few recommendations regarding the “*try-N-select-winner*” strategy:

**O1.** N should be as small as possible.
**O2.** Try to optimize a method before adding another one.
**O3.** The quality of the registration can be evaluated: i) By human-curation (although laborious, this is the most reliable method). ii) Automatic quantification of the quality of the registration relative to MEBRAINS. After masking the volumes with the MEBRAINS-mask, the following parameters can be evaluated: Pearson correlation; Normalized mutual information; SNR and peak-SNR; Mean Squared Error; Structural Similarity Index; Jaccard index; Dice Score; Hausdorff distance; Focal parameters for 3d images from the Image Quality Index toolbox (bias, correlation, divergency, entropy difference, root of mean squared error); Universal Image Quality Index (Vaiopoulos, 2011). All parameters should be normalized and scaled (0 – completely dissimilar; 1 - identical images), and can be calculated using MATLAB. The winner registration is established as the maximum value of the evaluated parameters, or of a metric defined on the space of all parameters (e.g., Euclidean distance).
c. *Run-N-select-high-probability-values*. The strategy works with volumes with discrete values such as atlases and involves the following steps:

1. Select N registration methods and run the registration of the same atlas (N ∼ 5).
2. Evaluate the quality of the registration and select M (M ≤ N) of the best registrations.
3. Build the probability distribution of values in corresponding voxels of the M selected volumes.
4. Build the resulting volume by giving to each voxel the value that has the occurrence probability greater than an optimal threshold. The optimal threshold depends on the overall probability distributions.
5. that higher N values are optimal. For example, we increased the number of registrations of the D99 atlas using both the registration of the D99-atlas to MEBRAINS and of the D99 atlas in NMT v2.0 space to MEBRAINS.

### Deep learning-based neuroimaging pipeline for automated processing of monkey brain MRI scans

Deep learning is becoming popular in the analysis of brain MR images, and is more widely used to MRI compared to other types of medical images (Zhao and Zhao, 2021). Deep learning has been used for pre-processing and analysing MR images, including brain segmentation, registration, noise reduction, resolution enhancement, restoration, and reconstruction (Zhao and Zhao, 2021). It has also been instrumental for computer-aided diagnosis, including lesion and tumor detection, and diagnostics of psychiatric and neurodegenerative disorders (e.g., Schizophrenia, Alzheimer’s disease, Parkinson’s disease, brain age estimation).

Traditional neuroimaging pipelines involve computationally intensive, time-consuming optimization steps, often requiring manual interventions (Henschel et al., 2020). To avoid these issues, we prepared two deep neural networks-based tools to work with the EBRAINS template:

#### U-Net Brain extraction tool for nonhuman primates (Wang et al., 2021)

This is a fast and stable U-Net based pipeline for brain extraction that exhibited superior performance compared to traditional approaches using a heterogenous, multisite non-human primate (NHP) dataset. The pipeline includes code for brain mask prediction (https://github.com/HumanBrainED/NHP-BrainExtraction.git), model-building, and model-updating, as well as macaque brain masks of PRIME-DE data (https://fcon_1000.projects.nitrc.org/indi/indiPRIME.html). A major advantage of the pipeline is that it uses a transfer-learning framework leveraging a large human imaging dataset to pre-train a convolutional neural network (U-Net Model), which is transferred to NHP data using a much smaller NHP training sample. Furthermore, the generalizability of the model can be improved by upgrading the transfer-learned model using additional training datasets from multiple research sites in the Primate Data-Exchange (PRIME-DE) consortium (136 macaque monkeys with skull-stripped masks repository, publicly available) (Milham et al., 2018).

We applied the package by carrying out these steps:

a. **Minimal pre-processing of the T1 images of the 10 monkeys included in the MEBRAINS template:**

- Conformed all images (FreeSurfer’s standard).
- Spatial adaptive non-local means filtering (using ANTS’s DenoiseImage).
- Bias field correction (using ANTS’s N4BiasFieldCorrection)
b. **Mask prediction** - use existing trained models to predict the mask for our data. The package provides 15 pre-trained models using different sets of data for transfer of learning and upgrading results. Each of the 15 models predicted a mask for each macaque anatomy including:

- 10 monkeys used to build MEBRAINS template, and 3 supplementary monkeys from our lab that will be included in later versions of the template.
- 21 monkeys from PRIME-DE (19 UC Davis and 2 U Minnesota). The goal of this process was to select the best performing models on our data.
c. **Supplementary model updating** - use the existing trained models and additional training datasets to improve the generalizability of the model:

- Select 7 models showing high performance in (b).
- Update each of these 7 models by supplementary training (40 epochs) using:

- Training data – 34 T1 images (10 used for MEBRAINS + 3 new from our lab; 21 from PRIME-DE (19 UC Davis and 2 U Minnesota)).
- Testing data: 66 T1 images (34 training data; 32 new data from KU Leuven). Fo all T1 images, ground-truth was derived from human-curated masks either created by us or taken form the repository from the U-net brain extraction package (https://fcon_1000.projects.nitrc.org/indi/indiPRIME.html, https://github.com/HumanBrainED/NHP-BrainExtraction.git).
d. Applications of the results:

- Use N-models to predict N versions of the mask for the same whole brain anatomy. N includes the 7 selected U-net models with their original parameters, and the 7 upgraded models (step c).
- Select the best result(s).
- If there was a clear winner, we used it. If there were more than one good approximations of the mask, we built a probability distribution for values (0 or 1) in each voxel. The final mask can be built by optimal thresholding of the probability distribution (“Run-N-select-high-probability-values” strategy).
- If necessary, adjust the result using manual adjustments and mathematical morphology applications in FSL, ANTS, AFNI and FreeSurfer

In all cases, the goodness of the predicted mask was evaluated by visual inspection or calculation of the dice score.

### Relative quality of the MEBRAINS template

To quantitatively evaluate the quality of our template relative to that of other templates, we used a method inspired by (Seidlitz et al., 2018). We chose for comparisons the following T1 templates: our MEBRAINS and ANTS10 templates, the NMT v2.0 (Seidlitz et al., 2018) and Yerkes19 (Donahue et al., 2018; Van Essen et al., 2012) templates, and the combination of the T1/T2 images of MEBRAINS and ANTS10. The two latter datasets were introduced to emphasize the usefulness of our multimodal approach. The processing of these 6 datasets included the following steps:

a. For each template, we segmented the amygdala (Am), caudate (Cd), claustrum (Cl), nucleus accumbens (NAc), putamen (Pu), white matter (WM), cortical gray matter (GM) and lateral ventricle (LV).
b. Normalization of the variations in T1 image intensity across each volume by N4 bias field correction (Avants et al., 2011) (using ANTS’s N4BiasFieldCorrection). T1/T2 images were generated from the original T1 and T2 images (without N4 bias field correction).
c. Using volume contraction (AFNI), we selected the kernel of each segment by excluding the external 3 voxels thick shell of each sub-cortical region.
d. We calculated the average gray matter (mean_GM_) of N randomly selected voxels (N = 50) for each segmented region (Am, Cd, Cl, NAc, Pu, and GM). For the white matter, we calculated the average white matter intensity (mean_WM_) of all voxels from the WM kernel. For LV, we calculated the standard deviation of the intensity of the cerebral spinal fluid (std_CSF_) of N randomly selected voxels. Both means and standard deviation included equal numbers of randomly selected voxels from the left and right hemisphere (N = 50). These values were used to calculate the following parameters, that represent contrast-to-noise (C2N) (Jang et al., 2022) and relative difference (KI):

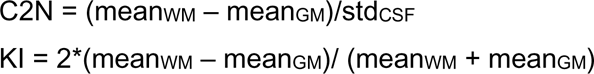
e. To evaluate the mean distribution of C2N and KI we performed the following steps:

**e1.** Compute the mean of C2N and KI by repeating their calculation 25 times, each time using a new set of 50 randomly selected voxels.
**e2.** Repeat step e1 2500 times to estimate the distribution of mean of the parameters.
**e3.** Steps e1-e2 were repeated for all 6 templates (the four T1 and the two T1/T2 datasets).
**e4.** Calculate the median values for each template and run a Kruskal-Wallis test followed by multiple comparison corrections.

## RESULTS

### MEBRAINS volumetric and surface templates

Our central goal was to build a population-based macaque brain template using multimodal imaging data to overcome limitations in the existing templates. Accordingly, we used MB to build three volumetric templates based on T1, T2 and CT brain images of 10 monkeys: MEBRAINS_T1 (Figure 3A), MEBRAINS_T2 (Figure 3B) and MEBRAINS_CT (Figure 4), respectively.

**Figure 3.**
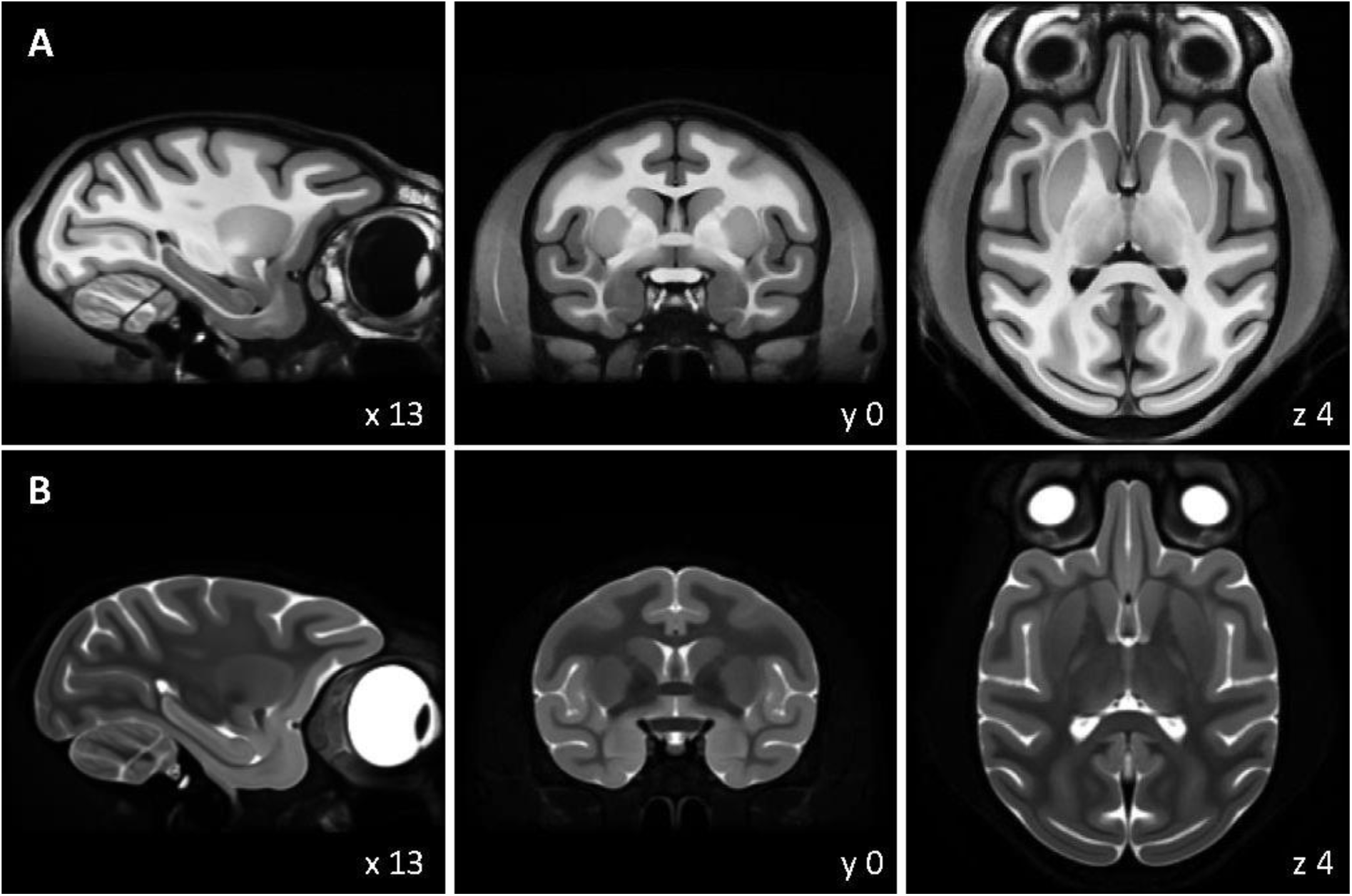
Three orthogonal sections of the MEBRAINS_T1 (A) and MEBRAINS_T2 (B) templates. The NIFTI-volumes used to create this figure can be found in supplementary material, and are also made publicly available via the EBRAINS platform from the Human Brain Project (https://doi.org/10.25493/5454-ZEA).

**Figure 4.**
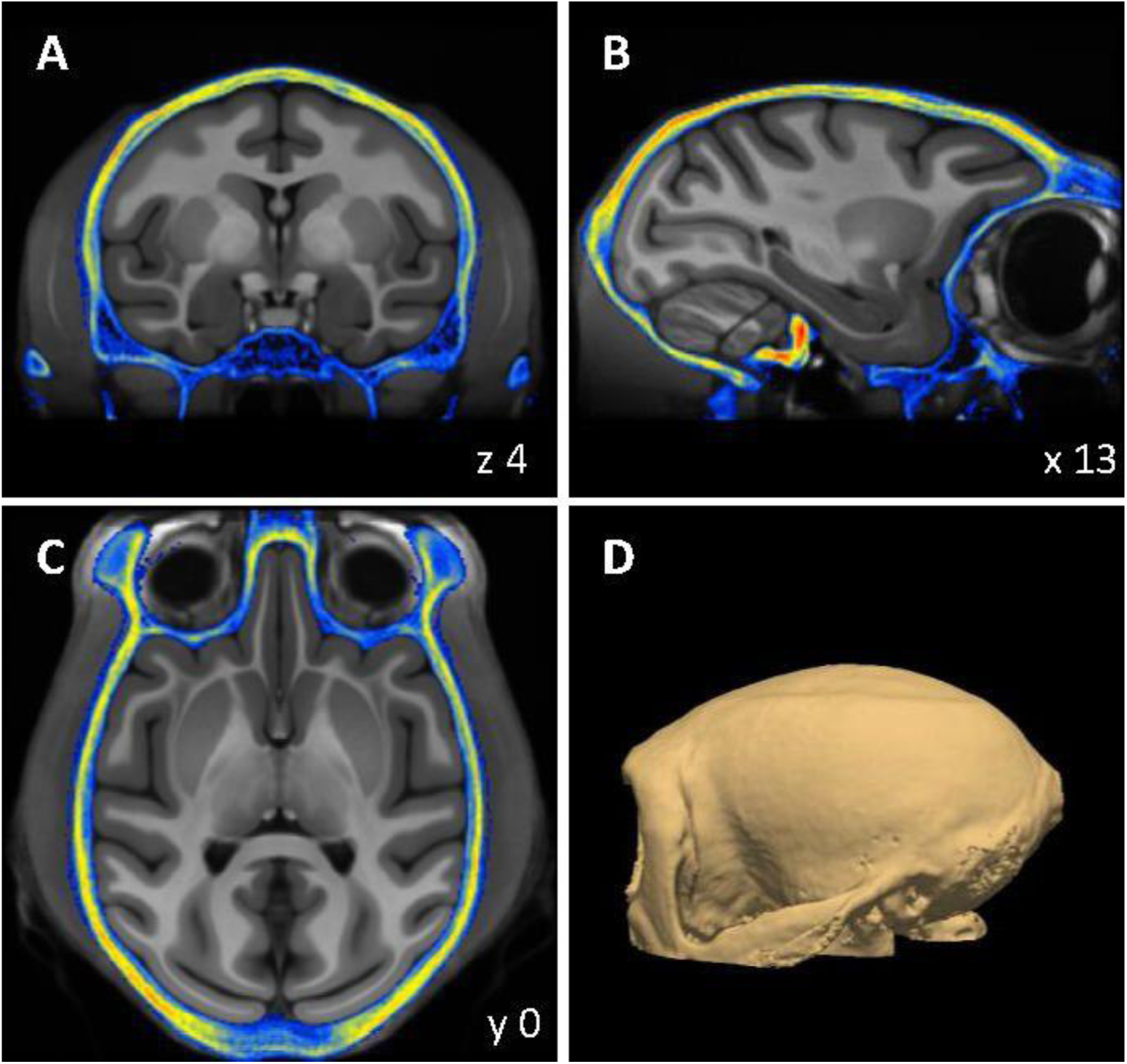
Three orthogonal sections (A-C) and 3D rendering (D) of the MEBRAINS_CT template. The corresponding NIFTI-volume can be found in the supplementary material, and is also made publicly available via the EBRAINS platform from the Human Brain Project (https://doi.org/10.25493/5454-ZEA).

Additionally, we created a second set of templates with the T1 and T2 brain images from the same 10 monkeys, but using ANTS, one of the few alternative tools besides MB that can rely both on T1 and T2 images for building templates (ANTS10_T1, Figure 5A and ANTS10_T2, Figure 5B). We found ANTS to result in a poorer tissue contrast compared to MB. Hence, we did not use it for our novel template, but to quantitatively compare the quality of the MEBRAINS templates relative to others.

**Figure 5.**
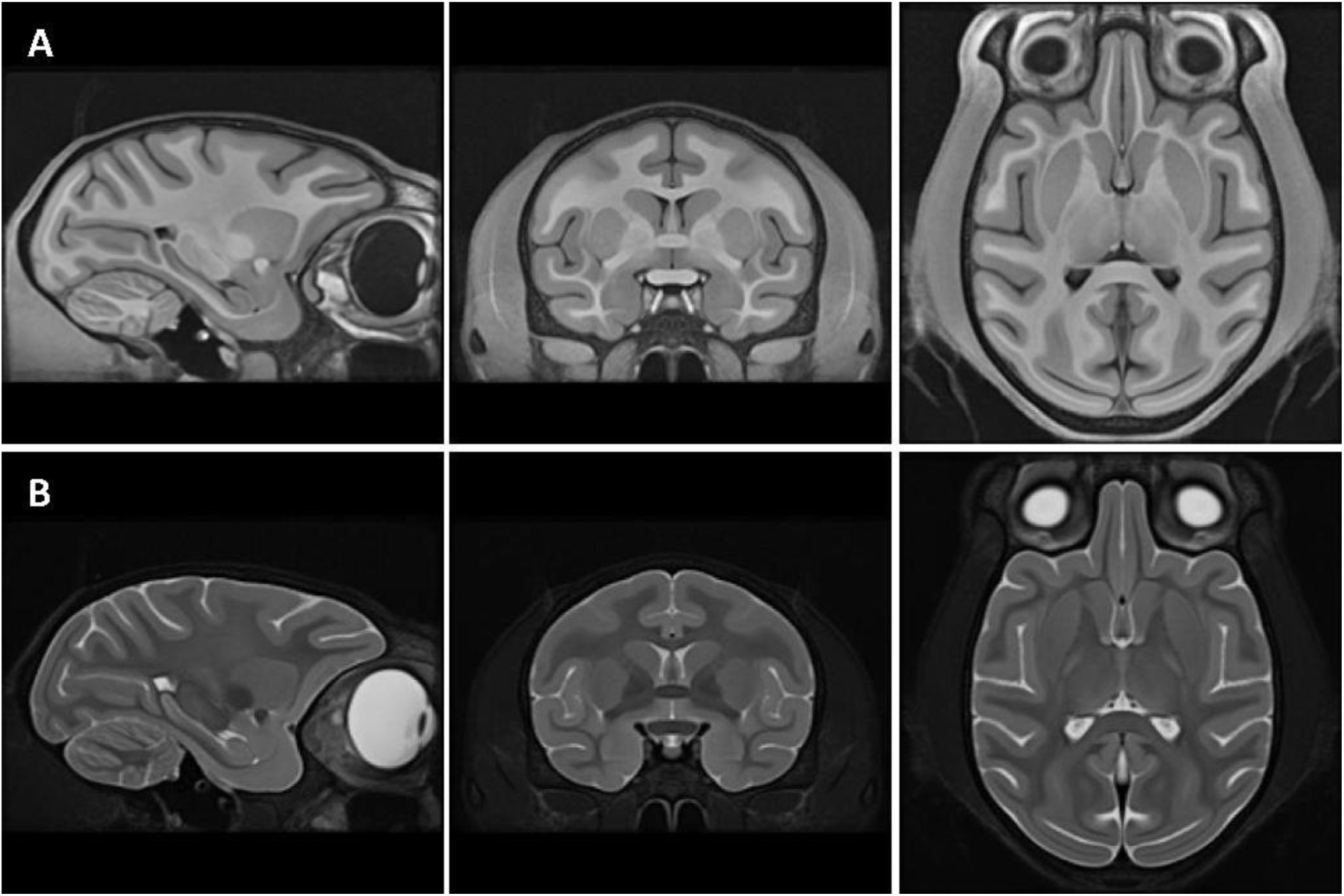
Three orthogonal sections of the ANTS10 templates generated from T1 (A) and T2 (B) images. To facilitate comparison with the corresponding MEBRAINS templates, the sections shown are the same as those depicted in Figure 1.

Finally, we also created a surface version of MEBRAINS, which will allow users to select between a folded or a flattened representation of the template’s cortex. We decided to use FreeSurfer to segment the white and grey matter of MEMBRAINS (Figure 6A), because it provided a better result than the grey and white matter masks generated by MB - as illustrated in Figure 6B, C. Note that, during the group-wise image registration process, MB generates tissue segmentations. Although the resulting probabilistic tissues do not necessarily correspond to anatomical parts of the brain, some of them provided a good approximation of the white and gray matter (Figure 6B, C). A supplementary merging and processing of the original MB-generated tissues may further improve the segmentation process. Yet given the satisfactory FreeSurfer results, we did not attempt this.

**Figure 6.**
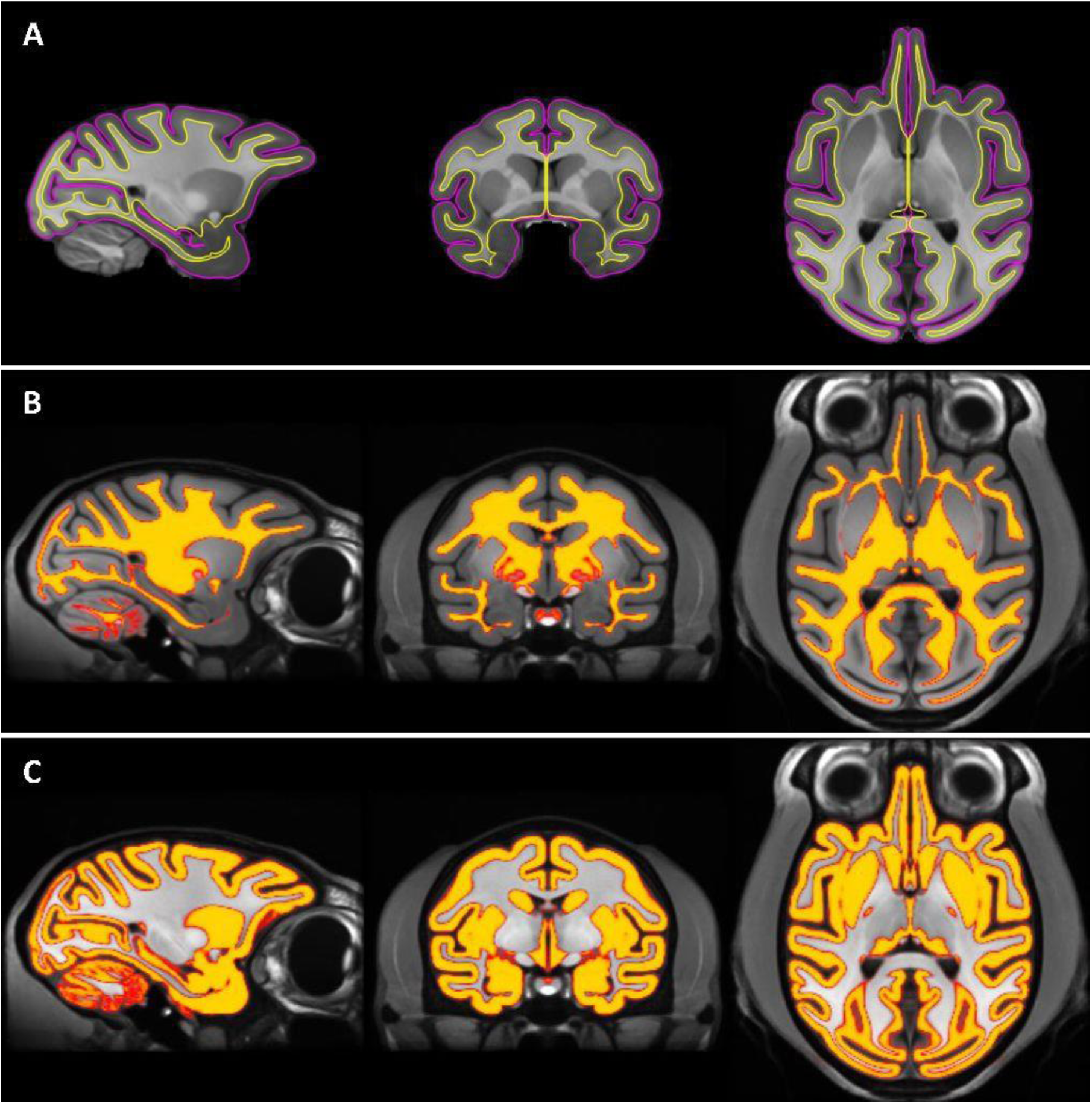
Generation of pial and white matter surfaces using FreeSurfer (A) and MB (B, C). (A) Pial (magenta) and white matter (yellow) boundaries overlaid on the MEBRAINS_T1 template. (B) White matter mask overlaid on the MEBRAINS_T1 template. (C) Gray matter mask overlaid on the MEBRAINS_T1 template. The sagittal, coronal, and horizontal sections depicted correspond to coordinates x13, y0 and z4, respectively.

### “Populating” the MEBRAINS template

It is essential for a template to be populated with neuroscience data. Indeed, a template becomes gradually more valuable by anchoring research results such as cyto-and myeloarchitectonic information, receptor distributions, task related activations, connectivity maps, electrophysiological data, and topographic maps such as retinotopic, somatotopic and tonotopic maps. In addition, it is important to link different template spaces. To start addressing these goals, we provided - in addition to white and grey matter segmentations based on FreeSurfer (Figure 6A) or MB (Figure 6B, C) - a human-curated segmentation of the anterior commissure and several major subcortical structures including the amygdala, nucleus accumbens, caudate, claustrum, putamen and pallidum (Figure 7A).

**Figure 7.**
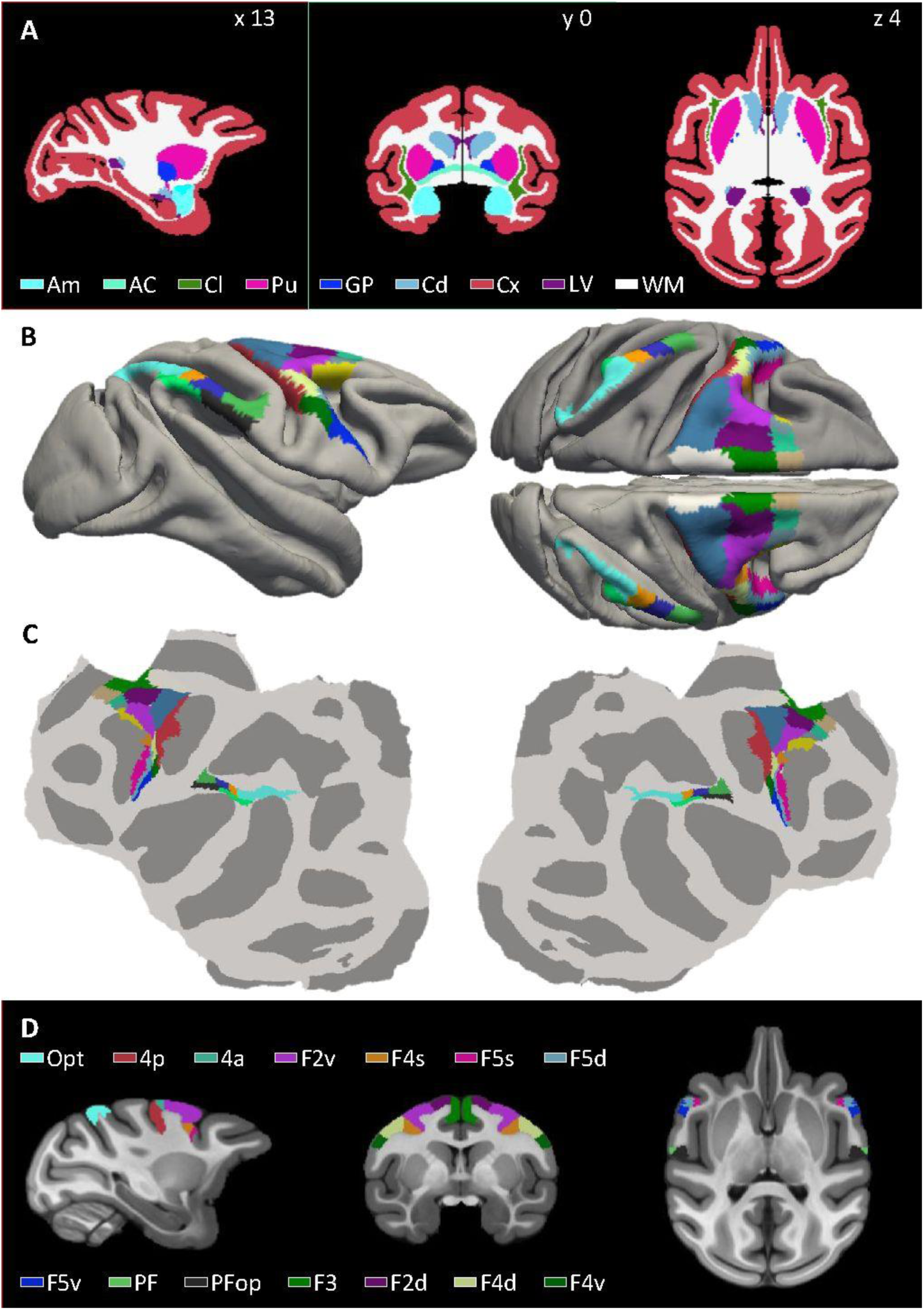
(A) Human curated segmentation of the cortical ribbon, white matter and lateral ventricles, as well as of diverse subcortical nuclei, and the anterior commissure. (B,C,D) Areas of the macaque inferior parietal lobule (Niu et al., 2021) and of the motor and pre-motor cortex (Rapan et al., 2021) warped from the Yerkes19 template to MEBRAINS. Areas are overlaid on the folded surface of MEBRAINS in (B), the flat maps in (C), and exemplary sections of MEBRAINS_T1 are shown in (D). Abbreviations: AC = anterior commissure; Am = Amygdala; CC = cerebral cortex; Cd=Caudate nucleus; Cl = Claustrum; GP = globus pallidus; LV = lateral ventricle; NAc = Nucleus accumbens; Pu=Putamen. The sagittal, coronal, and horizontal sections depicted in A and D correspond to coordinates x13, y0 and z4, respectively.

Furthermore, our recently published 3D cyto- and receptor architectonically-informed maps of the macaque monkey motor, premotor and parietal cortex were warped from YERKES19 space to the MEBRAINS surface template (Figure 7B), which were also represented on a cortical flat map (Figure 7C), and transformed into volumetric MEBRAINS space (Figure 7D). Since these areas were only available on the left hemisphere of the Yerkes19 template, and the MEBRAINS template is symmetrical, areas were mirrored to its right hemisphere.

### Registration of 3D datasets to MEBRAINS

The purpose of a template is to offer a standardized stereotaxic space for the analysis and/or visualization of neuroscience data, often requiring the co-registration of different volumes (e.g., individual brain anatomies, templates). Given the aforementioned advantages and limitations of MB, we propose a multi-method workflow with 4 major steps to integrate data into MEBRAINS space: Steps 1-3 encompass standardized pre- processing procedures, the actual computation of transformation functions (such as matrices and deformation volumes) necessary to register an anatomical image to MEBRAINS, as well as a quality assessments and improvements of the registration. Step 4 is only required if a data set instead of a structural anatomical volume needs to be registered, such as retinotopic maps, connectivity maps or parcellation schemes. In this case, steps 1-3 are performed with the reference anatomy, and the transformations/deformations functions are then applied to the associated datasets.

To demonstrate the validity and flexibility of our workflow, we first describe the result of our registration procedures when applied to some frequently used macaque brain templates, although they can be applied to any individual or averaged anatomical 3D volume. In a second step, we provide an example of how Step 4 can be implemented to transform a parcellation scheme of the macaque brain from the Yerkes19 surface to the MEBRAINS surface and volumetric templates.

#### Registration of other macaque brain templates to MEBRAINS

We considered the following macaque brain templates (Table 1; Figure 8): NMT v2.0, Yerkes19, D99, MNI macaque, F99, INIA19, ONPRC18 and 112RM-SL. Most of these templates are uni-modal (T1-weighted images) and skull-stripped, whereas MEBRAINS is a multi-modal (T1 and T2) template which includes the skull. Thus, these comparisons enabled us to test the aforementioned limitations of MB, and to demonstrate the usefulness of multi-method workflows for working with MEBRAINS. We used several methods (<u>“Try-N-select-winner” strategy, see methods</u>) from the library described in the methods (a – g; MB, ANTS, AFNI, MINC, ART, ITKsnap and FSL) to register the selected templates to MEBRAINS. MB performed well for the T1 templates in which the skull was not stripped (e.g., NMT v2.0), yet produced distorted registrations for many of the skull-stripped templates. The most optimal registration method for all registered templates was ANTS. Figure 8 shows ANTS10_T1, the 8 selected templates, and a meta-template (the average of the ANTS10_T1, and all template datasets, excluding 112RM-SL), all registered to MEBRAINS using ANTS. Furthermore, figure 8 also provides a unique opportunity to compare other templates with MEBRAINS. At qualitative level, MEBRAINS reveals comparable anatomical details as NMT V2.0, unlike the other templates.

**Figure 8.**
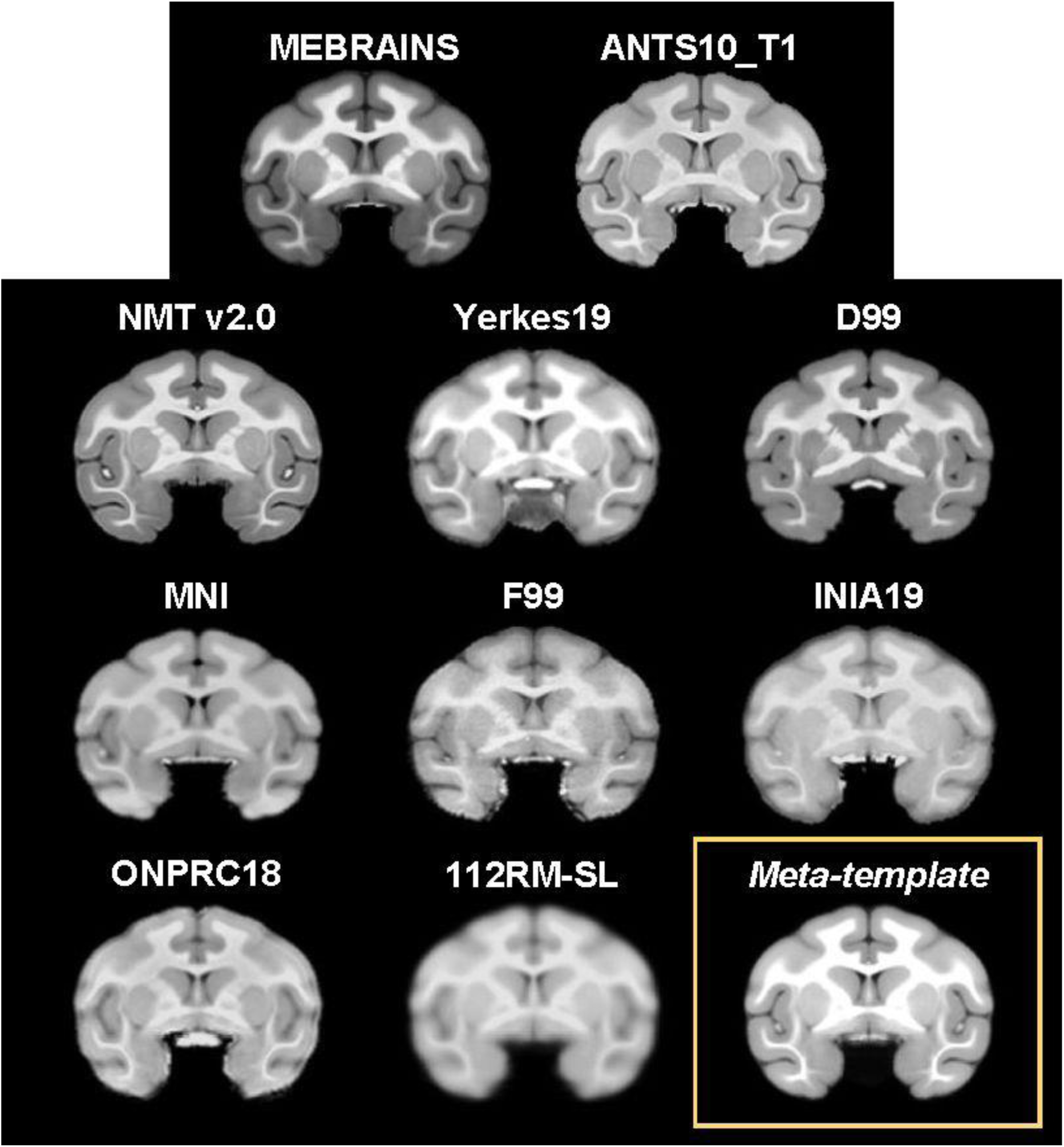
Eight of commonly used rhesus macaque brain templates (NMT v2.0 (Seidlitz et al., 2018), Yerkes19 (Donahue et al., 2018; Van Essen et al., 2012), D99 (Reveley et al., 2017), MNI (Frey et al., 2011), F99 (Van Essen, 2004), INIA19 (Rohlfing et al., 2012), ONPRC18 (Weiss et al., 2021) and 112RM-SL (McLaren et al., 2009)), as well as our ANTS10_T1 volume (i.e., the template built with ANTS using the same 10 datasets as MEBRAINS_T1) were registered to MEBRAINS using ANTS. The meta-template represents the average of all these datasets with the exception of 112RM-SL.

Figure 9 shows a quantitative evaluation of the quality of the registrations of the different templates with MEBRAINS (in Figure 8) using Pearson correlation and focal entropy differences -which was scaled to improve comparisons with the correlation method (0 – total dissimilarity; 1 – total similarity). Focal entropy was calculated for each coronal section using a symmetrical window radius of 7 voxels centered on each voxel and the results were averaged. Next, the differences between the average values for the registered and the reference (MEBRAINS) anatomies were calculated for each coronal section and averaged to obtain a value characterizing the entire volume. Both parameters provide an evaluation of how similar the compared anatomies are. Considering the range of values for both parameters (0.92-0.99), we conclude that all registrations have a good and relatively similar quality. The small individual variations also include differences between the intrinsic quality of the input image, which can be noticed by visual inspection in Figure 8).

**Figure 9.**
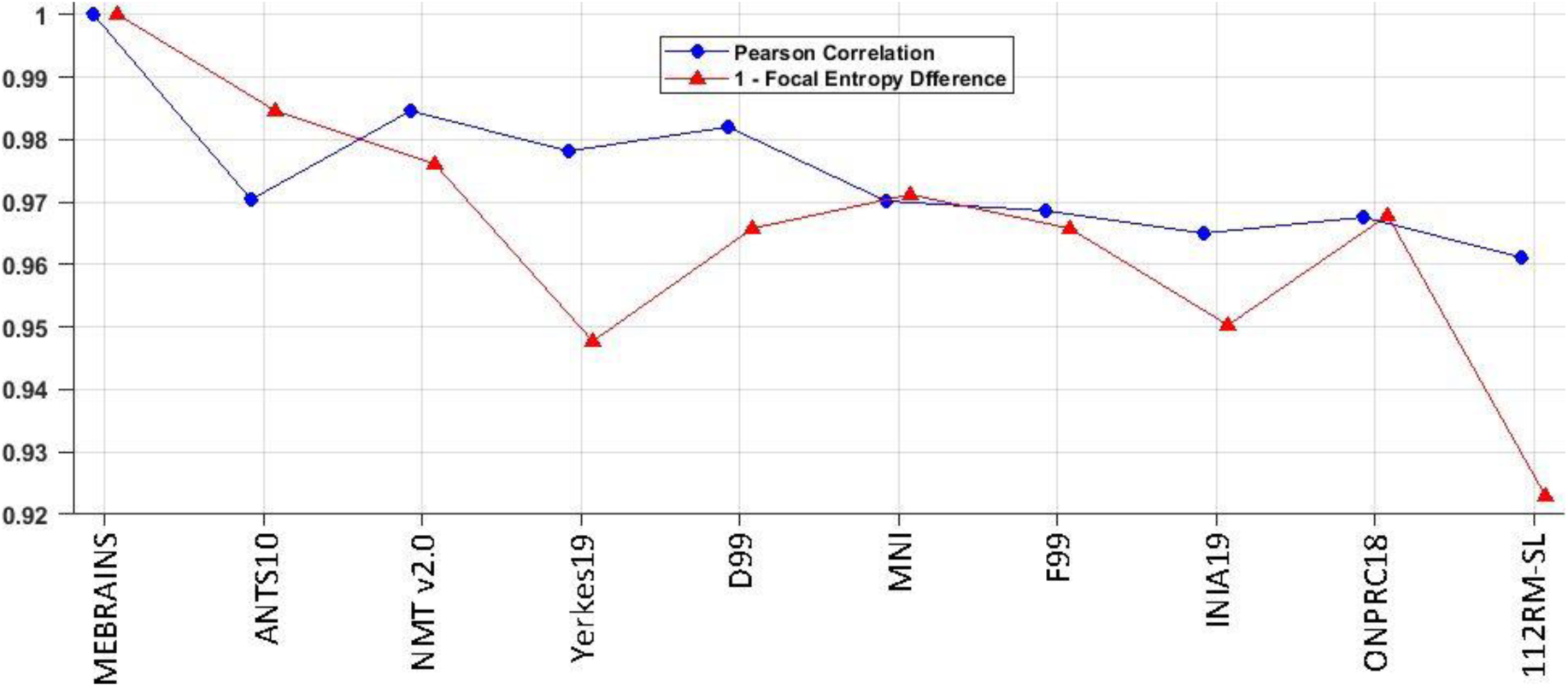
Pearson correlation and “1 – Focal Entropy Difference” (scaled to facilitate comparisons with the correlation method: 0 – total dissimilarity; 1 – total similarity) calculated for the reference anatomy MEBRAINS compared with the following templates: MEMRAINS, ANTS10_T1, NMT v2.0, Yerkes19, D99, MNI, F99, INIA19, ONPRC18 and 112RM-SL. Comparison of MEBRAINS with itself (value 1) provides the reference for the ideal registration.

#### Registration of a volumetric atlas to MEBRAINS

We here describe the result of the registration of the frequently used D99 atlas to MEBRAINS. We first registered the D99 template to MEBRAINS as described above using MB or ANTS and applied the “Try-N-select-winner” strategy (see methods). The resulting transformation objects (volume/matrix) were then applied to the associated D99 atlas using a nearest neighbourhood resampling algorithm (MB, Figure 10A; ANTS Figure 10B). Both registrations represent a good starting point for human-curated refinements.

**Figure 10.**
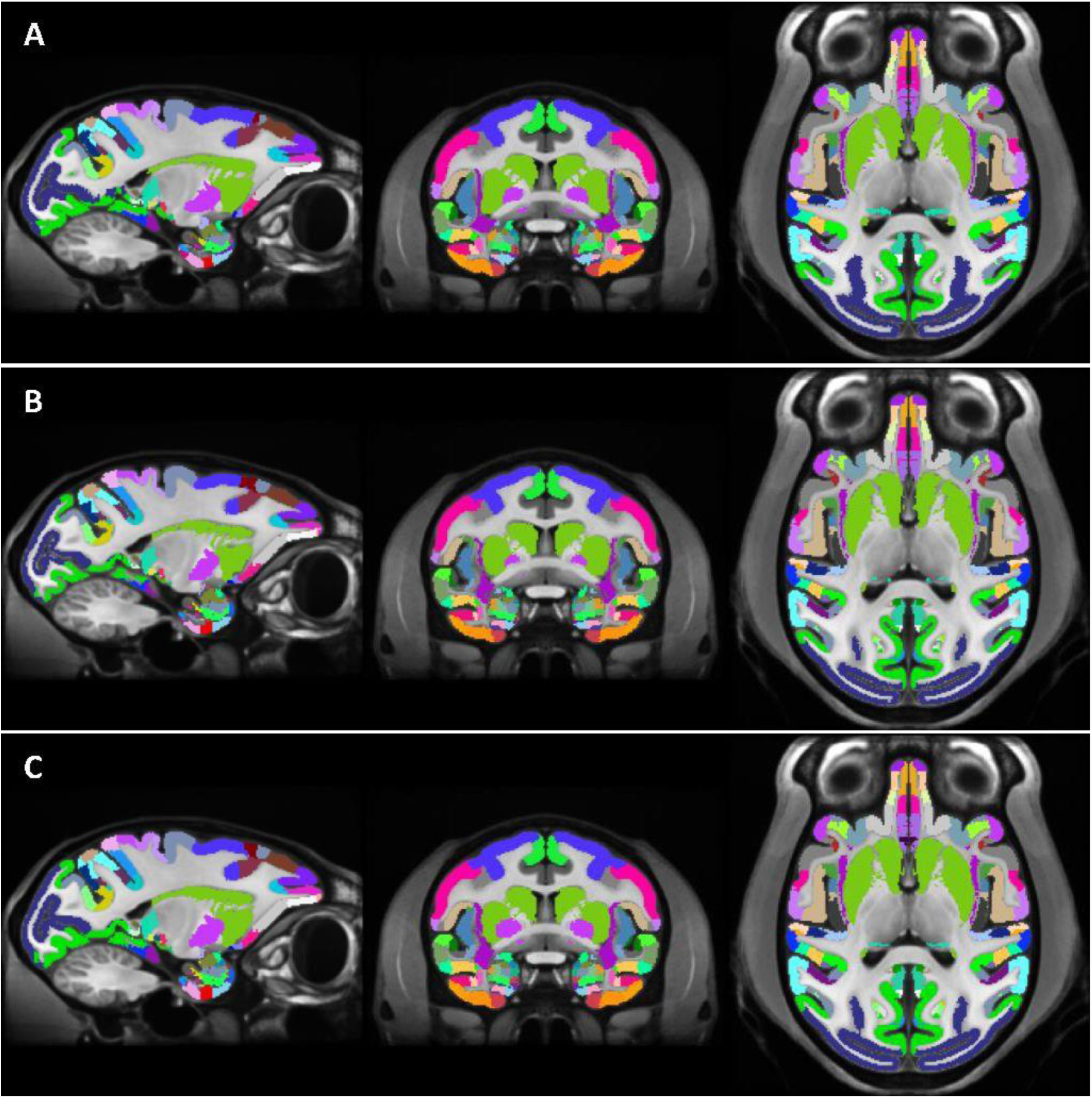
D99 atlas registered to MEBRAINS using the MB (A), ANTS (B) and “run-N-select-high-probability-values” (C) approaches. The different registrations of the atlas are overlaid on the MEBRAINS template.

We also performed the same registration (D99 atlas to MEBRAINS) using the “run-N- select-high-probability-values” strategy (Figure 10C). Because this method yields more information, given by the probability distribution of the voxel intensity values, than the single registration methods (Figure 10A, B), the resulting registration is more reliable.

#### Registration of a surface-based atlas to MEBRAINS

Since the 3D cyto- and receptor architectonically informed maps of the macaque motor, premotor and parietal cortex are associated with the Yerkes19 surface template, it was necessary to warp them to the MEBRAINS surface template using FreeSurfer, thereby establishing a link between both spaces. The ensuing labels can be visualized on the folded (Figure 7B) or flattened (Figure 7C) versions of the MEBRAINS surface template. Finally, they were transferred to the MEBRAINS volumetric template (Figure 7B).

### Deep learning-based neuroimaging pipeline for automated processing of monkey brain MRI scans

#### Automated brain extraction tool for nonhuman primates (U-NET) (Wang et al., 2021)

We performed supplementary training and updated the 7 existing models in the U-Net brain extraction package using 34 T1 images for training and 66 T1 images to test the mask prediction performances (see methods). The model training reached a dice score of 0.9882 ± 0.0005 (mean ± SEM) in epochs ranging between 35 to 39. The 7 upgraded models correctly predicted the mask in 85.71 ± 1.35 % (mean ± SEM) of the test brains and 94.96 ± 0.84 % of the trained brains. Moreover, more than one of the used models gave good predictions for the mask of the same brain. Accordingly, of 12 models used to predict the mask for each brain, 8.65 ± 0.27 (mean ± SEM) made good predictions for training and 7.97 ± 0.44 for testing data. Therefore, there is a substantial pool of good mask predictions for each brain allowing the use of either “*try-N-select-winner” or “run-N-select-high-probability-values”* strategies for brain extraction.

Figure 11 provides two example results of the winner for an ‘easy”, good quality anatomy, (Figure 11A) and for a more “difficult” lower quality anatomy (Figure 11C). As can be seen in Figure 11B, the dataset with the “difficult” anatomy requires longer training time than the “easy” anatomy before reaching the optimal solution. The example also emphasizes the robustness of the model, which is largely independent of the quality of the input data.

**Figure 11.**
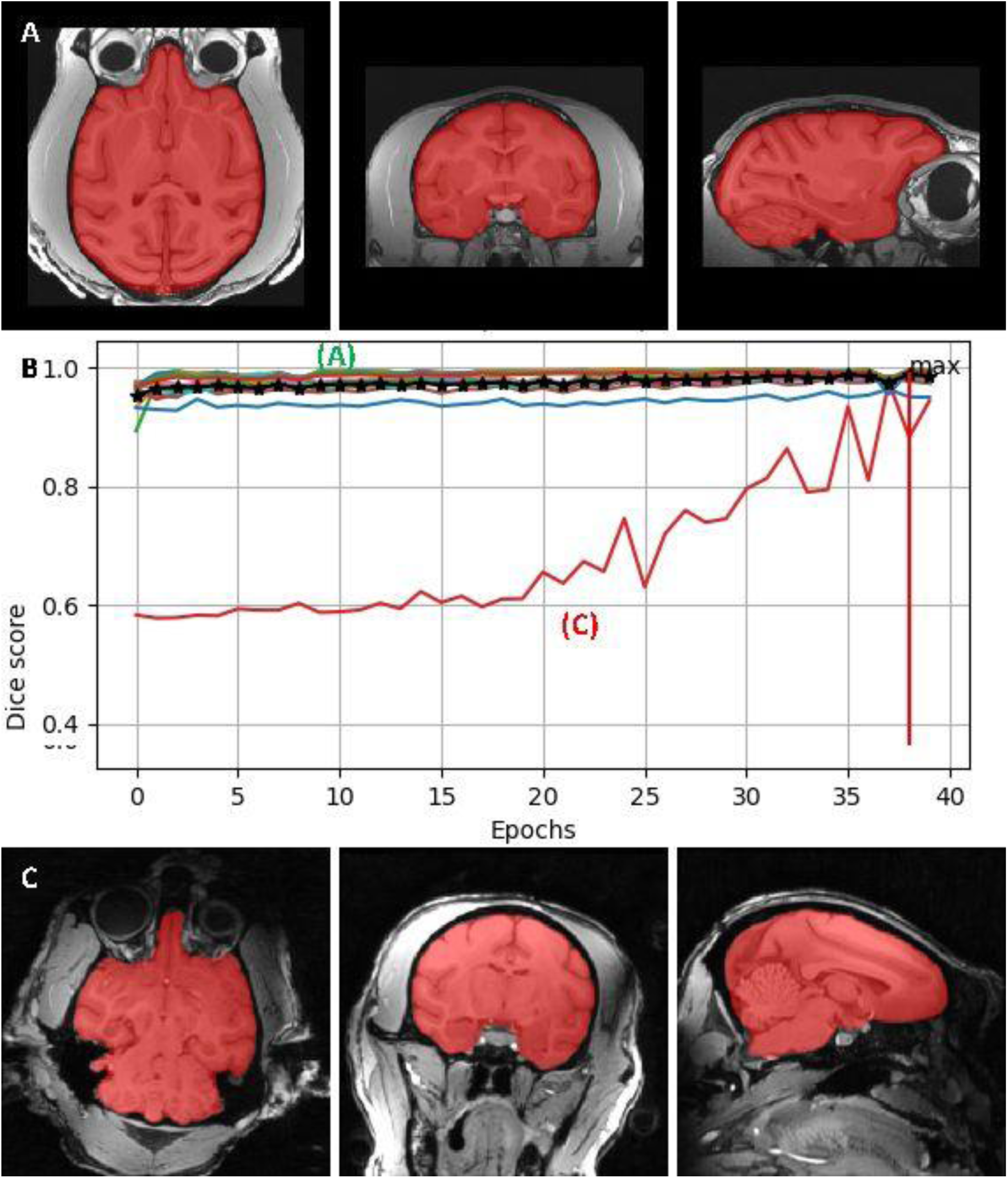
Masking performance of the U-net convolutional neural network using one example model. The predicted mask at the end of the training for an “easy” anatomy (A) and a “difficult” anatomy (C), and the dice score during the training (B). The performance for the “difficult” anatomy (red line in B) reached the optimal performance later than for the “easy” anatomy (green line in B). The maximum average dice score is 0.9887, and was reached in epoch 38.

### Relative quality of the MEBRAINS template

To quantitatively compare the quality of different templates, we segmented a number of anatomical structures from four T1 templates (MEBRAINS_T1, ANTS10_T1, NMT v2.0, Yerkes19) and two T1/T2 datasets (MEBRAINS_T1/T2, ANTS10_T1/T2) (Figures 12A). Depending on the quality of the template, the exact border of a structure may be difficult to estimate. Therefore, to be conservative in our comparison, we excluded the 3 most external voxels at each boundary of each of these compartments: for example, 3 voxels at the pial and 3 at the grey-white matter boundary for the cortical ribbon. As an example, Figure 12B shows the result of this process for MEBRAINS_T1. We used two different indices, inspired by (Seidlitz et al., 2018), to compare the quality of the templates (C2N and KI, see methods). The results presented in Figures 13 and 14 and Tables 2 and 3, support a few important conclusions regarding the possibility to distinguish different anatomical substructures of the brain in the different templates. First, the multi-modal templates MEBRAINS_T1/T2 and ANTS10_T1/T2 carry far more information compared to the unimodal ones. Hence, templates based on a combination of modalities allow improved segmentation of important brain structures. This is reflected in the larger C2N and KI values for the T1/T2 images. Notice that MEBRAINS_T1/T2 and ANTS10_T1/T2 (colored red and greed in Tables 2 and 3, respectively) outperform all other templates. Second, parameters for the T1-based templates show two different trends: C2N yields the largest values for the MEBRAINS_T1 template, while KI is dominated by NMT v2.0 (colored blue in Tables 2 and 3, respectively). Third, although NMT v2.0 is on par with the unimodal (T1) MEBRAINS, as shown by C2N and KI values, the multi-modal (T1/T2) approach in MEBRAINS provides a substantial advantage to all templates. Finally, comparison between MEBRAINS and ANTS10 demonstrates the superiority of MB compared to the ANTS for template generation.

**Figure 12.**
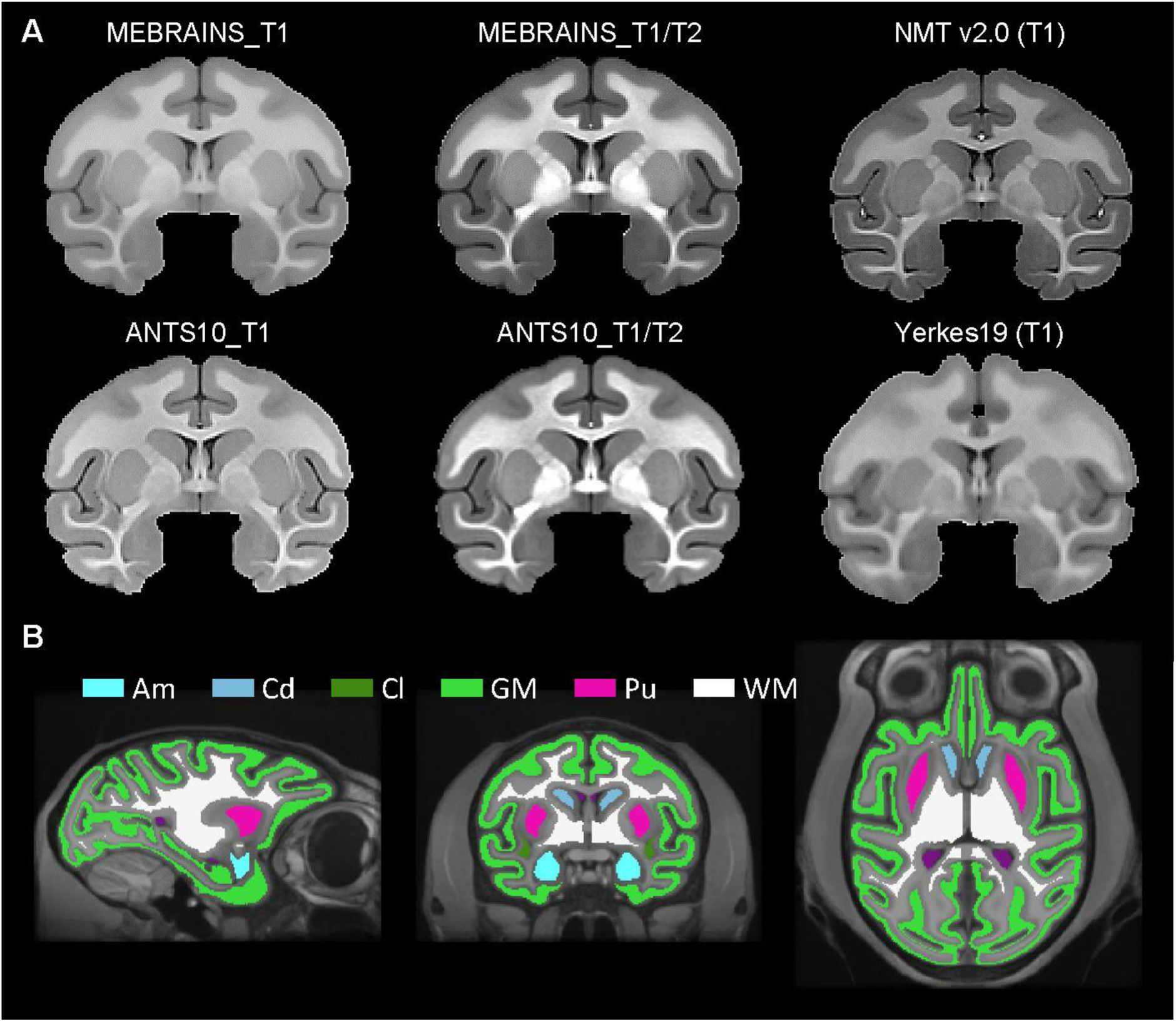
(A) Anatomies of the six templates used to quantitatively compare the quality of the EBRAINS template. (B) Structures that were selected for the MEBRAINS_T1 template: Am = Amygdala; Cd=Caudate; Cl = Claustrum; GM = cortical Gray Matter; Pu=Putamen; WM = White Matter.

**Figure 13.**
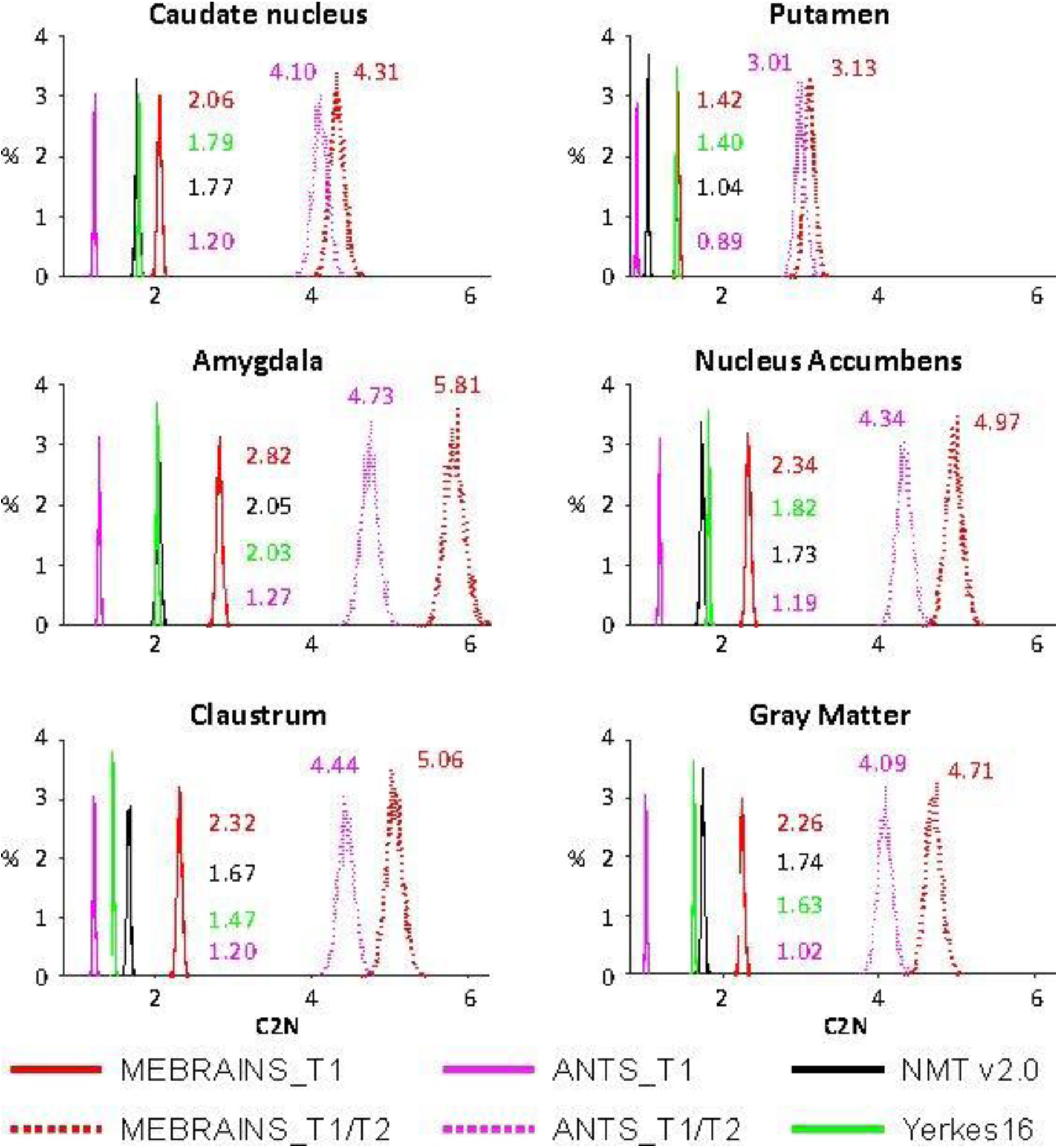
C2N parameter distribution of means for the templates shown in Table 2 and Figure 12A. Parameters were calculated for the 6 selected sub-structures separately, and numbers represent the median values.

**Figure 14.**
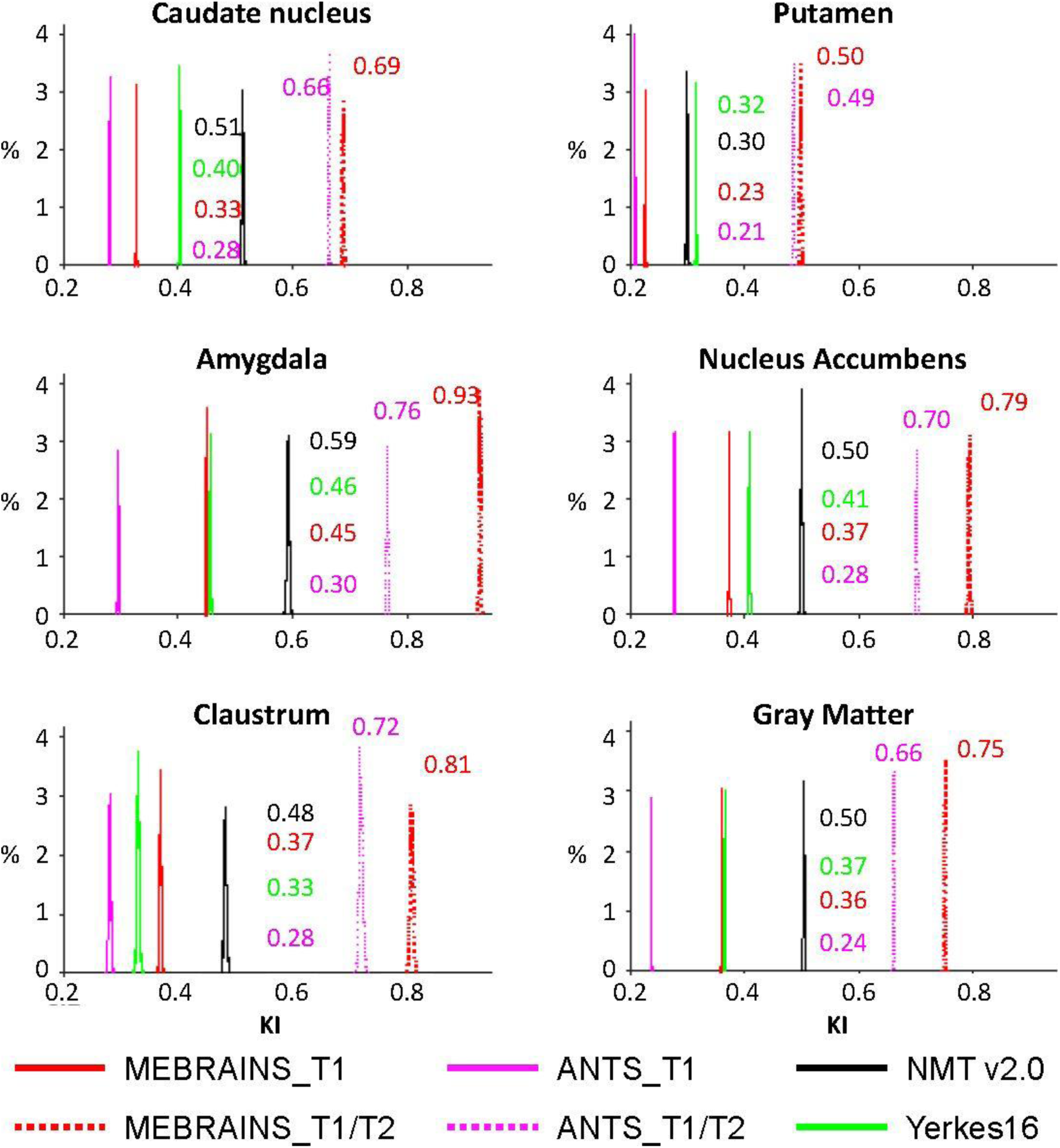
KI parameter distribution of means for the templates shown in Table 3 and Figure 12. Parameters were calculated for the 6 selected sub-structures separately, and numbers represent the median values.

**Table 2.** C2N median values for MEBRAINS_T1, MEBRAINS_T1/T2, ANTS10_T1, ANTS10_T1/T2, NMT v2.0 and Yerkes19. All pairs of medians are significantly different (p < 10^-8^) for each sub-structure. Fonts colored red, green (for T1/T2 images) and blue (for T1 images) outline the largest values of C2N. Abbreviations: Am = Amygdala; Cd = Caudate; Cl = Claustrum; NAc = Nucleus accumbens; Pu = Putamen; GM = cortical Gray-Matter.

| C2N | Cd | Pu | Am | NAc | Cl | GM |
| --- | --- | --- | --- | --- | --- | --- |
| **MEBRAINS_T1** | **2.06** | **1.42** | **2.82** | **2.34** | **2.32** | **2.26** |
| **MEBRAINS_T1/T2** | **4.31** | **3.13** | **5.81** | **4.97** | **5.06** | **4.71** |
| ANTS10_T1 | 1.20 | 0.89 | 1.27 | 1.19 | 1.20 | 1.02 |
| **ANTS10_T1/T2** | **4.10** | **3.01** | **4.73** | **4.34** | **4.44** | **4.09** |
| NMT v2.0 | 1.77 | 1.04 | 2.05 | 1.73 | 1.67 | 1.74 |
| Yerkes19 | 1.79 | 1.40 | 2.03 | 1.82 | 1.47 | 1.63 |

**Table 3.** KI median values. for MEBRAINS_T1, MEBRAINS_T1/T2, ANTS10_T1, ANTS10_T1/T2, NMT v2.0 and Yerkes19. All pairs of medians are significantly different (p < 10^-8^) for each sub-structure. Fonts colored red, green (for T1/T2 images) and blue (for T1 images) outline the largest values of KI. Abbreviations: Am = Amygdala; Cd = Caudate; Cl = Claustrum; NAc = Nucleus accumbens; Pu = Putamen; GM = cortical Gray-Matter

| KI | Cd | Pu | Am | NAc | Cl | GM |
| --- | --- | --- | --- | --- | --- | --- |
| MEBRAINS_T1 | 0.33 | 0.23 | 0.45 | 0.37 | 0.37 | 0.36 |
| **MEBRAINS_T1/T2** | **0.69** | **0.50** | **0.93** | **0.79** | **0.81** | **0.75** |
| ANTS10_T1 | 0.28 | 0.21 | 0.30 | 0.28 | 0.28 | 0.24 |
| **ANTS10_T1/T2** | **0.66** | **0.49** | **0.76** | **0.70** | **0.72** | **0.66** |
| **NMT v2.0** | **0.51** | 0.30 | **0.59** | **0.50** | **0.48** | **0.50** |
| Yerkes19 | 0.40 | **0.32** | 0.46 | 0.41 | 0.33 | 0.37 |

## DISCUSSION

We built a macaque brain template, MEBRAINS, in an attempt to mitigate common limitations of existing macaque templates. MEBRAINS is a multi-modal template that integrated relatively high resolution T1, T2 and CT modalities by using the MB toolbox (Brudfors et al., 2020). In addition, we developed both a volumetric and surface template. This approach will facilitate the combination of volumetric and surface data and enable the generation of flattened 2D maps of the cortex. As MEBRAINS is embedded in the EBRAINS environment which also houses human and rodent templates, and because other existing macaque templates have been registered to MEBRAINS, this will also expedite comparative research between macaques, humans and rodents.

To ensure the quality of both the data used to create MEBRAINS, and of the template itself, we applied a large spectrum of methods including those described in Marcus et al., 2013 (Marcus et al., 2013), tools borrowed from the image processing field tuned to evaluate image quality (e.g., see Figures 9, 11, 13, 14), and careful visual curation. Simple visual inspection of all the templates included in the present analysis (Figure 8) shows that the resolution and GM/WM contrast of MEBRAINS reveal a level of anatomical granularity and sharpness comparable to that of the NMT V2.0 template (Seidlitz et al., 2018), which is higher than that of most of the other templates, including the ANTS version of our template (ANTS10). This subjective impression was corroborated by the quantitative evaluation (Figures 13, 14), showing that the multi-modal MEBRAINS template represents anatomical details better than the other templates. The MEBRAINS_T1/T2 template presented the highest C2N values, indicating that the segmented structures have better signal to noise ratio compared to the other templates. Moreover, the multimodal character of MEBRAINS increases the discriminative power: MEBRAINS_T1/T2 yielded not only higher C2N (Jang et al., 2022) but also KI values compared to the remaining templates, including ANTS10_T1/T2. The latter finding is particularly interesting, because MEBRAINS and ANTS10 were constructed from the same 10 subjects. Specifically, this difference highlights the usefulness of multimodal approaches to construct brain templates.

Beyond the goal of creating a qualitative template, we adapted existing tools to register data to MEBRAINS (Figure 7, 10), to segment major anatomical structures (Figure 6, 7, 11, 12) and to generate surfaces (Figure 7). This included the adaptation of deep neural network approaches (U-NET), some of them also used in human research (FastServer) for processing monkey data.

Finally, we started to populate the MEBRAINS with previously published architectonic data (Donahue et al., 2016; Niu et al., 2020; Niu et al., 2021; Rapan et al., 2021). The comparison of such data with other parcellation schemes and future data sets will advance objective discussions about parcellations. In the future, we aim to refine the template by increasing the number of T1 and T2 images and by adding very high-resolution postmortem MRI anatomies. We also aim to register other functional data (e.g., probabilistic retinotopy data, category selective fMRI data, etc.) and increase the number of automatically segmented structures. Ultimately, we aim to obtain enough data to have a robust training set for our deep-learning based automated segmentation and registration of macaque data to MEBRAINS and any other template.

The MEBRAINS template represents the cornerstone of the “MEBRAINS Multilevel Macaque Brain Atlas” (https://atlases.ebrains.eu/viewer/monkey) developed in the framework of the Human Brain Project, which is freely available to the neuroscientific community via the interactive *siibra-explorer* on the EBRAINS platform (https://at-lases.ebrains.eu/viewer/monkey). Thus, MEBRAINS constitutes a spatial reference system to which a myriad of structural and functional *in vivo* and *post mortem* datasets with different degrees of spatial and temporal resolution will be anchored. Examples of *in vivo* datasets are electrophysiological, probabilistic retinotopy, category selective or resting state fMRI data as well as DTI datasets. *Post mortem* datasets include 3D-reconstructions of sections processed for visualization of cell bodies, myelinated fibres, neurotransmitter receptors distribution patterns or that of their subunits and/or the corresponding encoding genes, tractography datasets, as well as architectonic parcellation schemes of the macaque monkey brain. In this framework, the “Julich Brain Macaque Maps” (Donahue et al., 2016; Niu et al., 2020; Niu et al., 2021; Rapan et al., 2021), which are based on the quantitative analysis of differences in the distribution patterns of cell bodies and of multiple types of classical neurotransmitters, and to date had solely been available via the Yerkes19 surface template (Donahue et al., 2018; Van Essen et al., 2012), have now been registered to the MEBRAINS template. The maps and data associated with the MEBRAINS template can be used as entry point for higher level meta-analyses, or for guiding functional and interventional studies in MEBRAINS space. Furthermore, the richness of the EBRAINS meta-platform hosting the “MEBRAINS Multilevel Macaque Brain Atlas” and also representing humans and rodents in a unitary context enable efficient inter-species meta-analytical studies. Thus, MEBRAINS not only constitutes a technical improvement compared to previously published templates, but also facilitates cross-species comparisons.

In conclusion, via MEBRAINS we provide a novel population-based template of the macaque brain which was created using a multimodal approach and T1 and T2-weighted images. Quantitative evaluation of its quality demonstrated that it scores better than other unimodal templates. MEBRAINS constitutes the cornerstone of the “MEBRAINS Multilevel Macaque Brain Atlas” and has been populated with the cyto- and receptor-architectonically informed “Julich Brain Macaque Maps”. Importantly, MEBRAINS has been embedded in the framework of HBP’s EBRAINS platform, where it will enable the integration and analysis of multiple datasets of different spatio-temporal scales, and the comparison with other species.

## DATA AVAILABILITY

The volumetric and surface representation files of the MEBRAINS template are provided as supplementary files accompanying the manuscript and are also made freely available via the Human Brain Project platform EBRAINS (https://doi.org/10.25493/5454-ZEA).

## CODE AVAILABILITY

The following code is available on GitHub or software package webpages:

- Code used for creation of the templates is publicly available at (https://github.com/WTCN-computational-anatomy-group/mb). It requires the toolbox multi-brain for SPM12 and the commercial software MATLAB (Version R- 2018b). The repository includes example MATLAB scripts for template generation, registration to the template, different images co-registration
- FreeSurfer (https://surfer.nmr.mgh.harvard.edu/fswiki/DownloadAndInstall), ANTS (http://stnava.github.io/ANTs/), FSL (https://fsl.fmrib.ox.ac.uk/fsl/fslwiki/FslInstallation), AFNI (https://afni.nimh.nih.gov/), MINC (https://www.mcgill.ca/bic/software/minc), ART (https://www.nitric.org/projects/art/), Jip (http://www.nitrc.org/projects/jip), MRIcron (https://www.nitrc.org/projects/mricron), and ITK-SNAP (http://www.itksnap.org/pmwiki/pmwiki.php) are open source publicly available.
- U-Net Brain extraction tool for nonhuman primates (https://github.com/HumanBrainED/NHP-BrainExtraction) is publicly available and requires a python environment. Authors will provide by request the supplementary trained models.

## ACKNOWLEDGEMENTS

This work received funding from Fonds Wetenschappelijk Onderzoek-Vlaanderen (FWO-Flanders) G0D5817N, G0B8617N, G0C1920N, G0E0520N; the European Union’s Horizon 2020 Framework Programme for Research and Innovation under Grant Agreement No 945539 (Human Brain Project SGA3); KU Leuven C14/21/111, C3/21/027, IDN/20/016; the Federal Ministry of Education and Research (BMBF) under project number 01GQ1902.

We would also like to thank all members of the PRIMatE data sharing consortium (https://fcon_1000.projects.nitrc.org/indi/indiPRIME.html) for collecting, publishing and making their data available.

## AUTHOR CONTRIBUTIONS

1. Puiu F Balan – paper drafting, paradigm design, methods selection, data processing, data visualization
2. Qi Zhu – paper drafting, paradigm design, methods selection, data processing, data acquisition, data visualization
3. Xiaolian Li – data acquisition, data processing
4. Thomas Funck – paper drafting, data processing
5. Meiqi Niu – data processing, data visualization
6. Rapan – data processing, data visualization
7. Rembrandt Bakker – paper drafting, paradigm design, methods selection, data processing
8. Nicola Palomero-Gallagher – paper drafting, paradigm design, methods selection, project coordination, human-curated segmentation
9. Wim Vanduffel – paper drafting, paradigm design, methods selection, project coordination

